# Tunable kinetic destabilization governs RNA polymerase passage through a DNA-bound transcription factor

**DOI:** 10.64898/2026.07.29.741407

**Authors:** Noam Nago, Sergei Rudnizky, Adi Zohar, Nir Strugo, Hadeel Khamis, Carmit Burstein, Philippa Melamed, Ariel Kaplan

## Abstract

Eukaryotic transcription factors recognize short motifs, creating abundant binding sites within gene bodies and potential collisions with elongating RNA polymerases, yet how such encounters are resolved remains unclear. Here, we use optical tweezers to monitor RNA polymerase transcription through DNA-bound Egr-1, a zinc-finger transcription factor. Using DNA-fluctuation suppression as a readout of polymerase arrival, we show that Egr-1 delays elongation in an orientation- and rNTP-dependent manner, whereas force measurements indicate that RNAP does not bypass the TF by mechanical eviction. Instead, RNAP destabilizes the Egr-1–DNA complex over a short, structured interaction zone, increasing TF dissociation non-monotonically with distance. Monte Carlo simulations incorporating these kinetic changes recapitulate passage-time distributions. CpG methylation shortens Egr-1 residence time and largely eliminates the TF-dependent delay, suggesting a role for gene-body methylation in reducing kinetic barriers to elongation. These results reveal DNA-bound TFs as tunable barriers that locally shape transcription elongation.

## INTRODUCTION

Although transcriptional regulation is often discussed in terms of initiation, it is now clear that the elongation phase provides an additional and highly regulated layer of control across all domains of life^1–6^. The progression of elongating RNA polymerases is shaped by promoter-proximal pausing^7–11^, ubiquitous pauses^12^, sequence-specific pauses^13–15^, nascent RNA structures^16–18^, and direct interactions with regulatory factors^19^. In eukaryotes, transcription elongation is further regulated by chromatin structure, including nucleosomes^20–22^, chromatin remodelers, and histone chaperones^23^. By contrast, how other non-histone DNA-binding proteins modulate elongation is still poorly understood. This question is particularly relevant for eukaryotic transcription factors, which recognize short and degenerate DNA motifs, making potential binding sites abundant throughout the genome, including within gene bodies^24,25^. When such sites are occupied, elongating RNAPs must navigate DNA-bound proteins that may act as local barriers, potentially delaying elongation, promoting pausing or backtracking, or even inducing transcriptional arrest^26–28^. Because pausing during elongation regulates co-transcriptional processes such as RNA processing^29^ and alternative splicing^30,31^, these encounters have substantial regulatory potential. Yet the mechanisms by which RNAPs resolve encounters with DNA-bound TFs remain largely unknown.

Multi-subunit RNAPs, including bacterial and eukaryotic enzymes, translocate by a Brownian-ratchet mechanism^32,33^. During each nucleotide-addition cycle, RNAP fluctuates between pre- and post-translocated states, and incorporation of the incoming rNTP stabilizes the post-translocated state, thereby rectifying thermal motion into directional movement. Obstacles on the DNA track can interfere with this cycle by increasing the time RNAP spends in the pre-translocated state, thus enhancing the likelihood of backtracking^34–36^. Recovery from the backtracked state requires either diffusional realignment^37,38^ or transcript cleavage, which is stimulated by elongation factors such as TFIIS in eukaryotes and GreA/GreB in bacteria^34,35^. Recent work showed that GreA can facilitate passage of bacterial RNAP through a strongly bound obstacle^39^, but whether elongation factors are required to overcome smaller, moderately bound barriers is unclear.

The early growth response protein 1, Egr-1, also known as Zif268, provides a well-suited model for studying RNAP– TF encounters. Egr-1 is induced by diverse stimuli, including growth factors^40^, neurotransmitters^41^, hormones^42,43^, and stress^44^, and binds DNA through three zinc fingers (ZFs), each contacting 3 bp of DNA^45,46^.

Previous studies have established a 9-bp GCGTGGGCG consensus binding motif^47^. However, most genomic binding sites deviate from this consensus, and many of these deviations are evolutionarily conserved^48^, suggesting functional importance. *In vitro*, such sequence variation alters Egr-1 affinity and binding kinetics^48–50^, but even substantially diverged sites retain appreciable affinity^49^, suggesting that they may be occupied at physiological Egr-1 concentrations^51,52^. These properties make Egr-1 a useful model for testing how elongating RNAPs navigate sequence-specific DNA-bound proteins that may occur within transcribed regions.

Traditional biochemical and structural approaches have provided valuable insights into transcription elongation, but they cannot directly follow the dynamic interactions between RNAP and DNA-bound proteins in real time. Here, we used single-molecule optical tweezers to monitor RNAP passage through DNA-bound Egr-1 and independently probe the mechanical and kinetic properties of the encounter. Our results show that a complex and tunable interplay between TF kinetic stability and RNAP dynamics controls passage.

## RESULTS

### Non-perturbative detection of RNAP elongation

To monitor elongation by individual RNAP molecules, we designed a DNA construct containing the T7A1 bacterial promoter followed by a 27-bp T-less cassette. An *E. coli* RNAP initiation complex was assembled on this template and allowed to transcribe into the cassette before stalling by omission of UTP from the reaction buffer. Elongation-competent complexes were enriched and ligated to an alignment segment connected to double-stranded DNA handles bearing either biotin or digoxigenin modifications (Fig. S1a,b, Methods). The resulting DNA molecules were tethered between two optically trapped beads. To localize RNAP, we gradually unzipped the DNA by increasing the trap separation and then relaxed the force to allow rezipping (Fig. 1a). The presence of RNAP was detected as a localized force increase in the unzipping trace (Fig. 1b, gray), consistent with previous observations for RNAP^53^ and other DNA-binding proteins^26,48,54–58^. To verify that the stalled complexes remained elongation-competent after enrichment and tethering, we transferred the tethered construct into a separate laminar-flow channel containing all four rNTPs at 500 µM and performed repeated unzipping cycles (Fig. 1b, green). In successive cycles, the RNAP-dependent force barrier appeared at progressively smaller numbers of base pairs unzipped, indicating that RNAP had moved downstream along the template between measurements. This time-dependent shift in barrier position (Fig. 1c) confirmed that the stalled complexes retained the ability to resume transcription after rNTP addition.

**Figure 1:**
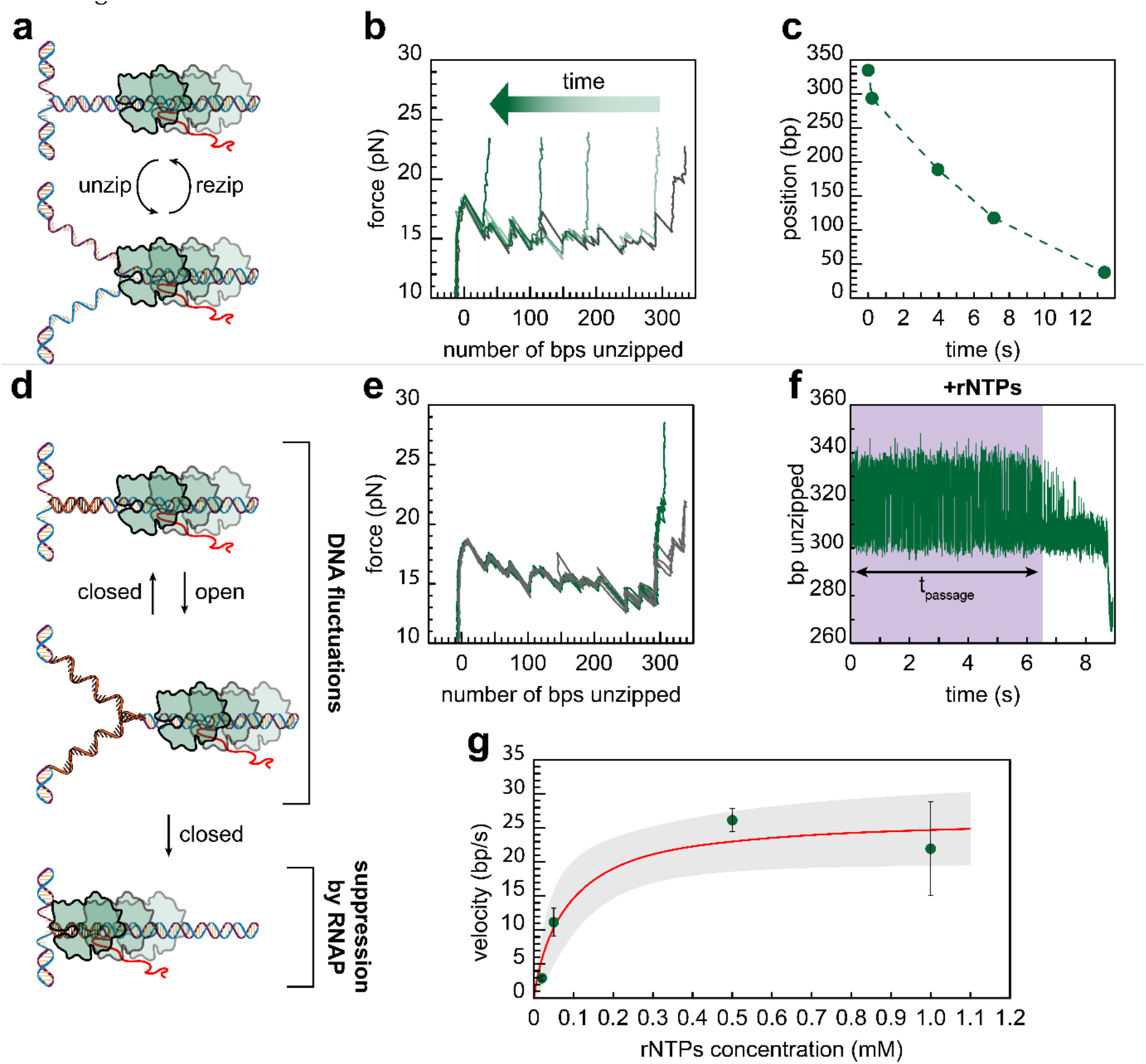
Non-perturbative measurements of RNAP translocation on the DNA. **a)** Schematic of the repetitive unzipping assay. A single stalled RNAP elongation complex is tethered between two optically trapped beads, and the DNA is repeatedly unzipped until RNAP is detected as a localized increase in force. **b)** Representative repetitive unzipping traces from the same RNAP complex. Before addition of rNTPs, RNAP is detected at approximately 350 bp unzipped position (gray). After addition of rNTPs, RNAP advances along the template and is detected at progressively smaller unzipping positions in successive cycles (green traces). **c)** RNAP position as a function of time for the experiment shown in **b. d)** Schematic of the DNA-fluctuation assay. A stalled RNAP elongation complex is localized by unzipping, after which the DNA is partially unzipped to a predetermined position where thermal opening and closing of the DNA fork generate spontaneous fluctuations. The tether is then transferred to a laminar-flow channel containing rNTPs to restart transcription. RNAP translocation into the fluctuating region suppresses DNA fluctuations, providing a label-free readout of polymerase arrival. **e-g)** Representative fluctuation-based measurement. RNAP is first localized by unzipping in the absence of rNTPs (**e**, gray line). The DNA is then held at fixed trap positions and transferred to rNTP-containing buffer, where RNAP arrival suppresses DNA fluctuations (**f**). Finally, the tether is returned to rNTP-free buffer and unzipped again to determine the final RNAP position (**e**, green line). **g)** RNAP elongation velocity as a function of rNTP concentration, calculated as the transcribed distance divided by the measured passage time. Data are mean ± s.e.m, n = 27, 6, 38, 6.

We previously demonstrated that thermal fluctuations in a partially unzipped DNA fork can report on the dynamics of protein-DNA interactions^50,57^, as binding within the fluctuating region suppresses the fluctuations and dissociation leads to their reappearance. We adapted this method to monitor RNAP elongation (Fig. 1d). After locating RNAP in rNTP-free buffer (Fig. 1e, gray), we positioned the unzipping fork approximately 60 bp downstream of the RNAP active site, held the trap positions fixed, and monitored spontaneous fluctuations within a ~20-bp region of the DNA (Fig. 1f). The construct was then transferred into an rNTP-containing channel to restart transcription. Arrival of RNAP at the fluctuating region suppressed the fluctuations, allowing us to define the passage time as the interval between entry into the rNTP-containing channel and fluctuation suppression (Fig. 1f, purple shading). To confirm RNAP translocation, we returned the construct to rNTP-free buffer and performed an additional unzipping cycle, which revealed a ~40-bp shift in RNAP position relative to its initial location (Fig. 1e, green). Notably, in this assay the transcribing polymerase is effectively shielded from external force by the fluctuating fork, and no tags or labels are required, so potential perturbations to RNAP activity are minimized. We calculated elongation velocity by dividing the transcribed distance by the measured passage time. The velocity showed the expected hyperbolic dependence on rNTP concentration (Fig. 1g), saturated above 0.1 mM rNTPs, and was consistent with earlier work^56^. We therefore used 0.5 mM rNTPs in all subsequent experiments.

### A bound Egr-1 delays elongation in an orientation- and rNTP-dependent manner

To determine how a DNA-bound transcription factor affects elongating RNAP, we introduced a consensus Egr-1 binding site 22 bp downstream of the active site of the stalled RNAP (Fig. 2a). Because the Egr-1 binding site is asymmetric, its orientation determines which zinc finger RNAP encounters first. In this initial construct, RNAP encounters zinc finger 1 first, followed by zinc fingers 2 and 3; we therefore refer to this as the ZF123 orientation. We then measured RNAP passage through this region in the presence of 69 nM Egr-1 DNA-binding domain (Fig. 2b), a concentration well above the reported *K*_*D*_^50^, such that the binding site is expected to be occupied most of the time. Despite the absence of accessory elongation factors, RNAP was able to transcribe through the Egr-1-bound site and reach the fluctuating region. However, passage was delayed in the presence of Egr-1 (Fig. 2c,d, blue). Thus, a bound Egr-1 does not form an absolute block to elongation but imposes a measurable kinetic barrier that RNAP can overcome.

**Figure 2:**
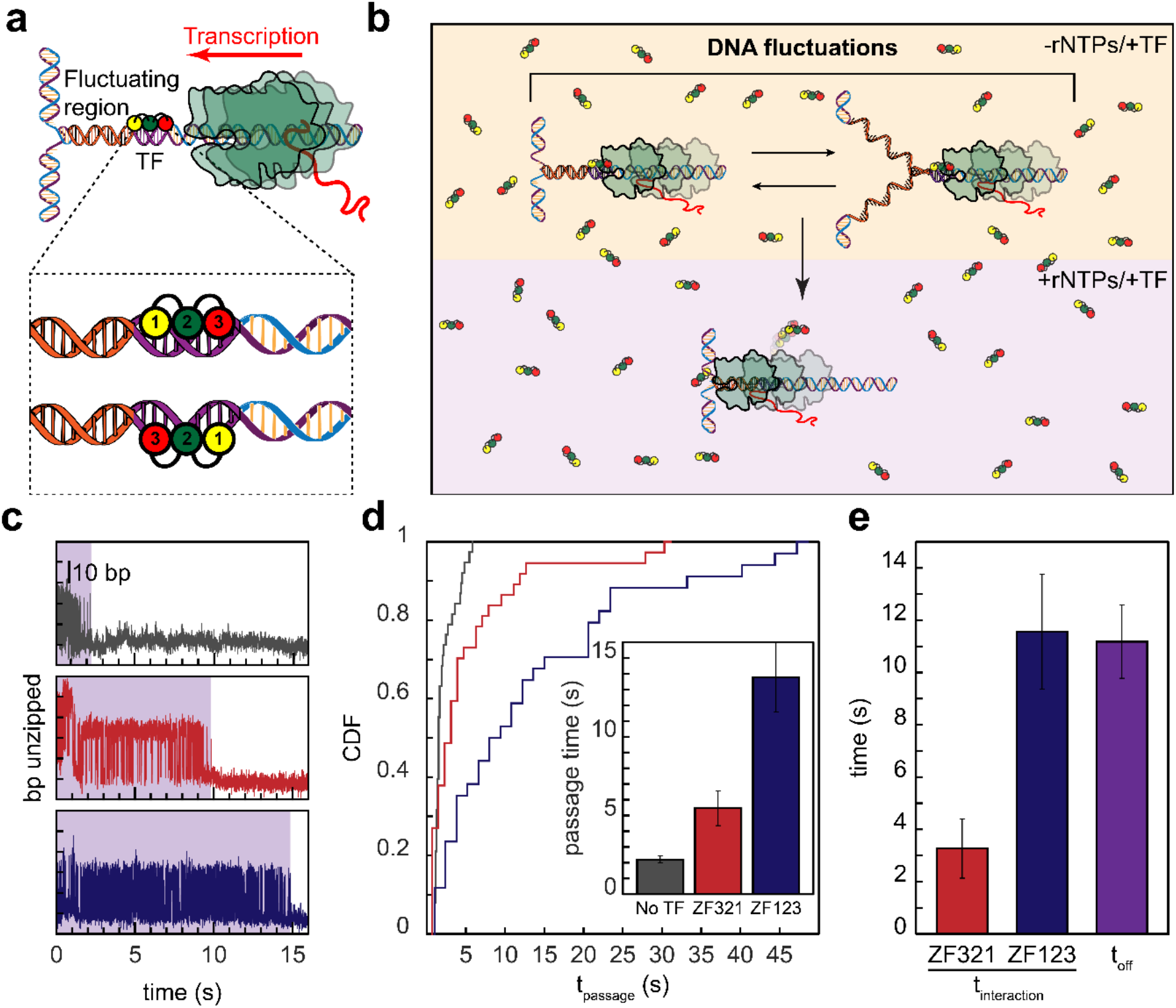
Egr-1 delays RNAP passage in an orientation-dependent manner. **a)** Schematic of DNA constructs containing a stalled RNAP elongation complex and an Egr-1 binding site positioned 22 bp downstream of the RNAP active site. The binding site was inserted in two opposite orientations, such that RNAP encounters the Egr-1 zinc fingers in either the ZF123 or ZF321 orientation. **b)** A single stalled RNAP complex is tethered and unzipped to reach the fluctuation region in a medium containing 69 nM Egr-1, and then transferred to a medium containing both Egr-1 and rNTPs (0.5 mM). **c)** Representative traces showing DNA fluctuations before and after RNAP arrival in the absence of Egr-1 and in the presence of Egr-1 bound in the two orientations. The purple shading indicates the passage time. **d)** Cumulative distributions of RNAP passage times in the absence of Egr-1 and in the presence of Egr-1 in the ZF321 and ZF123 orientations. Inset, mean passage times for the same conditions. Data are mean ± s.e.m.; n = 38, 37, and 34 molecules for no Egr-1, ZF321, and ZF123, respectively. Statistical comparisons are reported in Table S1. **e)** RNAP–Egr-1 interaction times for the ZF321 and ZF123 orientations, calculated for each orientation as the difference between the mean passage time measured in the presence of Egr-1 and the mean passage time measured in its absence. Error bars represent propagated s.e.m. from the corresponding passage-time measurements.

Because the measured passage time includes both elongation along the DNA and the additional time required to overcome bound Egr-1, we calculated the “interaction time” by subtracting the mean passage time measured in the absence of Egr-1 from the passage times measured in its presence (Fig. 2e). This interaction time was comparable to the separately measured mean residence time of Egr-1 on DNA (Fig. S2), raising the possibility of a simple passive mechanism, in which RNAP stalls at the bound TF and waits until Egr-1 dissociates spontaneously. We therefore tested two predictions of this passive waiting model. First, although the asymmetric Egr-1 binding site imposes a defined orientation on the protein, a passive model would predict that passage time should not depend on the orientation of the bound TF. Hence, we generated a second construct in which the binding site was reversed, so that RNAP encounters the protein in the opposite orientation, ZF321. RNAP was again delayed by a bound Egr-1 (Fig. 2c,d, red), but the delay was significantly shorter than in the ZF123 orientation. Thus, passage depends on the orientation of Egr-1 relative to the direction of transcription, suggesting that RNAP does not simply wait for spontaneous TF dissociation.

Second, we repeated the passage measurements at limiting rNTP concentration (Fig. S3). In a passive waiting model, the interaction time should primarily reflect spontaneous Egr-1 dissociation and should therefore be largely independent of rNTP concentration. For the ZF321 orientation, the interaction time was short relative to the total passage time, preventing us from reliably resolving any rNTP dependence. In contrast, the interaction time in the ZF123 orientation increased substantially under limiting rNTP conditions. Together with the orientation dependence, this result supports a mechanism in which RNAP does not merely wait for Egr-1 to dissociate but instead modulates the encounter in an rNTP-dependent manner. One possible explanation is that RNAP mechanically destabilizes or displaces the bound TF. We therefore next asked whether RNAP can generate sufficient force to rupture the Egr-1–DNA complex.

### RNAP-generated forces are insufficient to mechanically displace a bound Egr-1

To test whether RNAP could overcome Egr-1 by mechanical displacement, we directly compared the forces generated by RNAP with the forces required to rupture the Egr-1–DNA complex. We first measured Egr-1 rupture forces by unzipping DNA through the bound TF. Our previous work showed that the mechanical stability of Egr-1 is asymmetric^48^: the force required to remove Egr-1 by DNA unzipping depends on the direction in which force is applied. Measurements under the present experimental conditions confirmed this asymmetry, with significantly higher rupture forces in the ZF123 orientation than in the ZF321 orientation (Fig. 3a,b). For comparison, we measured the force required to remove a stalled RNAP from DNA by unzipping through the complex (Fig. 3c,d). The RNAP rupture force was lower than the rupture force of Egr-1 in the ZF123 orientation, but significantly higher than that of Egr-1 in the ZF321 orientation (Fig. 3g,h). Thus, the mechanical stability of a stalled RNAP–DNA complex falls between the two orientation-dependent Egr-1–DNA rupture forces. Finally, we measured the maximum force generated by an actively elongating RNAP, its stall force, by allowing the polymerase to transcribe against a static DNA fork (Fig. 3e,f).

**Figure 3:**
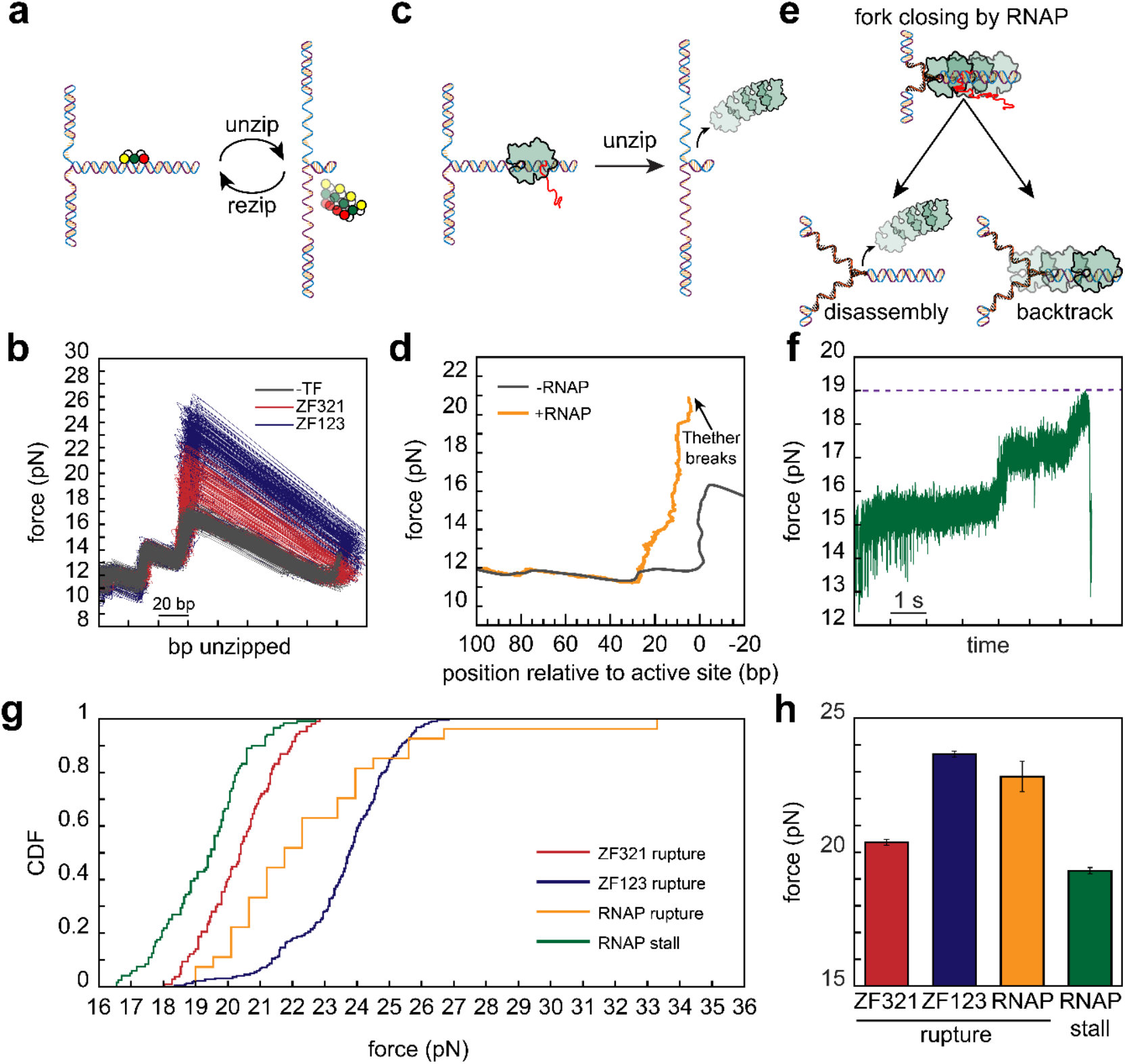
RNAP-generated forces are insufficient to mechanically remove Egr-1. **a)** Schematic of the Egr-1 rupture-force assay. A DNA molecule containing an Egr-1 binding site is unzipped until the bound TF is removed from the DNA. **b)** Representative unzipping traces showing rupture of the Egr-1–DNA complex in the ZF123 and ZF321 orientations. Gray traces show DNA unzipping in the absence of Egr-1. **c)** Schematic of the RNAP rupture-force assay. A DNA molecule containing a stalled RNAP elongation complex is unzipped until RNAP is removed from the DNA. **d)** Representative unzipping trace showing RNAP rupture from DNA, compared with DNA unzipping in the absence of RNAP. **e)** Schematic of the RNAP stall-force assay. After RNAP reaches the fluctuating fork, continued transcription closes additional segments of the unzipped DNA, increasing tether tension until RNAP stalls and subsequently backtracks or dissociates from the template. The stall force was defined as the maximum force measured during stalling. **f)** Representative force trace showing RNAP arrival at the fluctuating region and the measured stall force. The dashed magenta line marks the maximum force. **g)** Cumulative distributions of Egr-1 rupture forces, RNAP rupture forces, and RNAP stall forces. n_Egr1, 321_=107, n_Egr1, 123_=205, n_RNAP, break_=27, n_RNAP, stall_=119. Statistical comparisons are reported in Table S2. **h)** Corresponding mean forces from the data shown in **g**, shown as mean ± s.e.m.

These measurements used the same geometry as the passage assay, but acquisition was continued after RNAP reached the fluctuating fork. As RNAP advanced, it progressively closed the fork and increased the force on the DNA tether until forward translocation ceased. The polymerase subsequently backtracked or dissociated from the template (Fig. 3e,f; Fig. S4a–c). We defined the RNAP stall force as the maximum force reached before backtracking or dissociation. Consistent with previous measurements^56^, the RNAP stall force was lower than the force required to mechanically remove Egr-1, especially in the more stable ZF123 orientation (Fig. 3g,h; Fig. S4d). This force asymmetry is consistent with the orientation-dependent differences in passage time. However, the magnitude of the RNAP stall force indicates that RNAP cannot overcome Egr-1 through a simple force-driven eviction mechanism in which the polymerase directly pushes the TF off the DNA, suggesting a more complex passage mechanism in which RNAP may alter the kinetic stability of the Egr-1–DNA complex.

### RNAP destabilizes Egr-1 in a non-monotonic, distance-dependent manner

Given that RNAP-generated forces were insufficient to explain mechanical eviction of Egr-1, we next asked whether RNAP instead alters the kinetic stability of the Egr-1–DNA complex by modulating Egr-1 association and dissociation rates. During active elongation, however, the distance between RNAP and the TF binding site changes continuously, making it difficult to assign individual binding or dissociation events to a defined RNAP–TF separation. We therefore performed static measurements using stalled RNAP complexes, in which Egr-1 binding kinetics could be measured at defined distances from the RNAP active site (Fig. S5a).

We constructed DNA templates in which the proximal edge of the Egr-1 binding site was positioned at distances ranging from 12 to 58 bp downstream of the RNAP active site (Fig. 4a), and characterized the ZF321 orientation. Because Egr-1 binding kinetics are sensitive to sequence context flanking the binding site^50^, each construct was expected to have its own intrinsic association and dissociation rates. We therefore measured Egr-1 kinetics for each construct both in the presence and absence of RNAP, allowing RNAP-induced changes to be quantified relative to a matched sequence context. These constructs were introduced into buffer containing 69 nM Egr-1 and repeatedly unzipped to confirm both RNAP position and Egr-1 binding (Fig. 4b). Next, the unzipping fork was held near the Egr-1 binding site, allowing TF binding and dissociation events to be detected as changes in the DNA fluctuation pattern (Fig. 4c; Fig. S5b-i). Egr-1 association and dissociation rates were extracted from the distributions of unbound- and bound-state lifetimes (Fig. 4d,e; Fig. S5j,k), respectively, and compared with rates measured on the same construct in the absence of RNAP.

**Figure 4:**
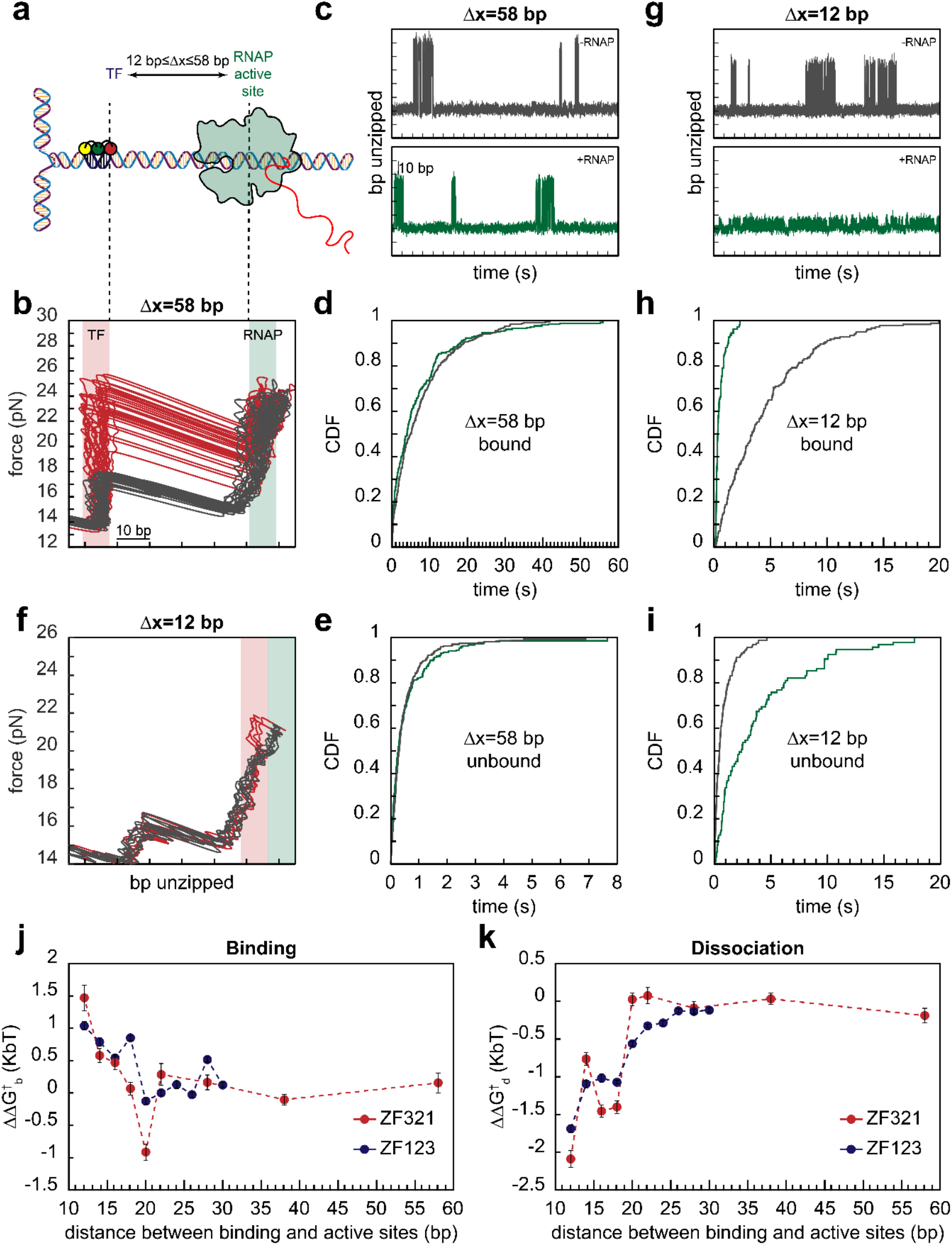
RNAP destabilizes Egr-1 binding to DNA in a non-monotonic, distance-dependent manner. **a)** Schematic of DNA constructs containing a stalled RNAP elongation complex at a fixed position and an Egr-1 binding site positioned with its proximal edge 12–58 bp downstream of the RNAP active site. **b)** Representative repetitive unzipping trace confirming the positions of stalled RNAP and Egr-1 for a construct in which the Egr-1 binding site is located 58 bp downstream of the RNAP active site. The Egr-1-bound trace is shown in red, and a reference unzipping trace of the same DNA molecule in the absence of Egr-1 is shown in gray. **c)** Representative DNA-fluctuation traces reporting Egr-1 binding and dissociation events in the presence (green) or absence (gray) of RNAP for the 58-bp construct. **d)** Cumulative distributions of bound-state lifetimes in the presence (green) and absence (gray) of RNAP for the 58-bp construct, from which dissociation rates, k_off_, were extracted. **e)** Cumulative distributions of unbound-state lifetimes in the presence (green) and absence (gray) of RNAP for the 58-bp construct, from which association rates, k_on_, were extracted. **f)** Representative repetitive unzipping trace confirming the positions of stalled RNAP and Egr-1 for a construct in which the Egr-1 binding site is located 12 bp downstream of the RNAP active site. **g)** Representative DNA-fluctuation traces reporting Egr-1 binding and dissociation events in the presence (green) or absence (gray) of RNAP for the 12-bp construct. **h)** Cumulative distributions of bound-state lifetimes in the presence (green) and absence (gray) of RNAP for the 12-bp construct, from which dissociation rates, k_off_, were extracted. **i)** Cumulative distributions of unbound-state lifetimes in the presence (green) and absence (gray) of RNAP for the 12-bp construct, from which association rates, k_on_, were extracted. **j)** RNAP-induced changes in the apparent transition-state energy for Egr-1 association, shown for the ZF123 and ZF321 orientations. **k)** RNAP-induced changes in the apparent transition-state energy for Egr-1 dissociation, shown for the ZF123 and ZF321 orientations. Energy changes were calculated from the measured rates, and error bars represent propagated errors from the corresponding rate estimates.

Given that elongation complexes contact downstream DNA over approximately 14–18 bp from the active site^59,60^, we initially expected Egr-1 to be sterically excluded from binding sites within this region. Surprisingly, Egr-1 binding was still detectable at sites inside this nominal footprint (Fig. 4f). Although close proximity to RNAP reduced the amplitude of the observed DNA fluctuations (Fig. 4g), binding and dissociation events could still be reliably detected, allowing both association and dissociation rates to be extracted (Fig. 4h-i, Fig. S5j,k).

The presence of RNAP significantly increased the Egr-1 dissociation rate and decreased its association rate, but only when the binding site was located within ~20 bp of the RNAP active site (Fig. S5j,k). A similar trend was observed in experiments in the ZF123 orientation (Fig. S6). To quantify these RNAP-induced kinetic changes in energetic terms, we converted the measured association and dissociation rates into apparent transition-state energy differences, ΔΔ*G*^‡^, for binding and dissociation, respectively (Fig. 4j,k). For each Δ*x* construct, ΔΔ*G*^‡^was calculated relative to the rate measured for the same DNA sequence in the absence of RNAP, thereby isolating the effect of RNAP proximity from sequence-dependent differences in intrinsic Egr-1 binding kinetics. This analysis revealed a non-monotonic dependence on RNAP–TF distance, suggesting that specific structural features of RNAP, rather than generic steric proximity alone, alter the kinetic barriers for Egr-1 binding and dissociation.

### Passage is governed by an orientation-dependent interplay between TF destabilization and RNAP backtracking

Together, these results suggest a passage mechanism in which RNAP does not simply collide with Egr-1 and wait for spontaneous TF dissociation. Rather, as RNAP approaches the bound TF, they mutually affect one another: Egr-1 hinders RNAP progression, and RNAP destabilizes the Egr-1–DNA complex. Both effects depend on RNAP–TF distance. Because such encounters can also induce RNAP backtracking, successful passage should depend not only on TF dissociation, but also on the balance between forward translocation, backtrack entry, and recovery from the backtracked state.

If this mechanism is correct, the distance-dependent destabilization measured in the static experiments, together with the intrinsic kinetic properties of RNAP extracted from passage experiments on naked DNA, should be sufficient to quantitatively explain RNAP passage in the presence of Egr-1. To test this, we performed stochastic Monte Carlo simulations of transcription through an Egr-1-bound template (Fig. 5a). The simulations incorporated the experimentally measured transition-state energy changes as a function of RNAP–TF distance (Fig. 4j,k), while RNAP elongation and backtracking parameters were constrained by the naked-DNA passage experiments (Fig. 5b). This kinetic-destabilization model successfully reproduced not only the mean passage times, but also the full passage-time distributions for ZF321 (Fig. 5c), supporting the idea that RNAP promotes passage by increasing the dissociation rate of a bound Egr-1 at specific distances. In contrast, a passive model, in which RNAP translocates until it encounters Egr-1 and then waits for spontaneous TF dissociation, failed to reproduce the experimentally observed passage times.

**Figure 5:**
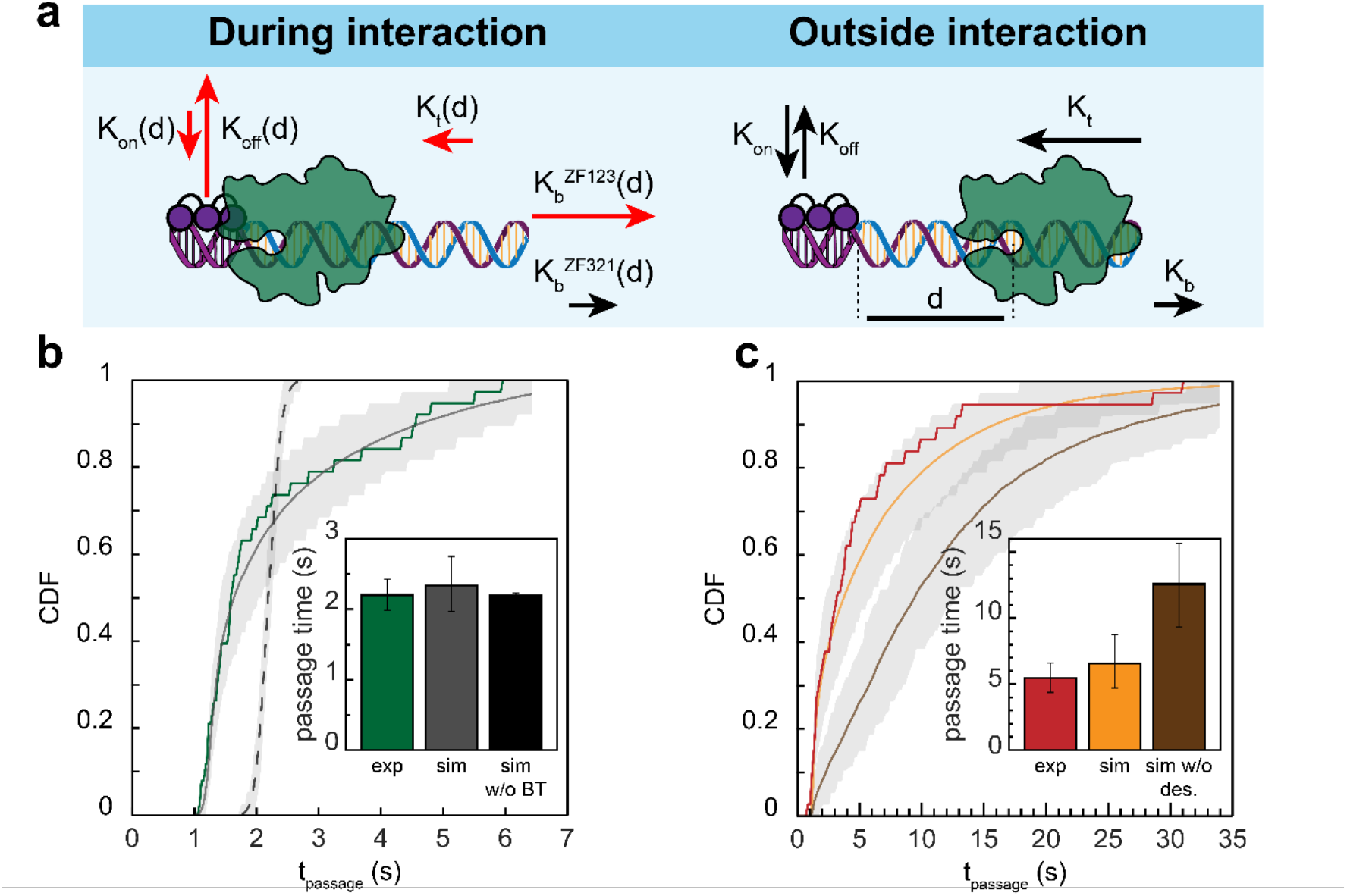
Interplay between Egr-1 destabilization and RNAP dynamics governs passage.. **a)** Model for RNAP passage through a DNA-bound TF. Outside the interaction region, RNAP elongation and Egr-1 binding kinetics are independent. Within this region, RNAP progression and Egr-1 binding kinetics become coupled: Egr-1 can delay RNAP elongation, whereas RNAP proximity can alter Egr-1 association and dissociation rates. **b)** Stochastic Monte Carlo simulation of RNAP passage in the absence of Egr-1. The solid dark gray line shows the simulated passage-time distribution obtained by fitting the experimental data to determine RNAP elongation and backtracking parameters. The green line shows the distribution of passage times in the absence of Egr-1 (same data as in Fig. 2d). The dashed gray line shows the fit of a model without backtracking. The gray band represents the simulation interval containing 90% of simulated datasets. Inset, corresponding mean passage times. Experimental data are shown as mean ± s.e.m.; simulation error bars indicate the central 90% interval of the distribution of replicate means. **c)** Stochastic Monte Carlo simulation of RNAP passage in the presence of Egr-1 in the ZF321 orientation. The orange line shows simulations using RNAP elongation parameters obtained from passage on naked DNA in panel b, and the position-dependent transition-state energy for Egr-1 association and dissociation measured in Fig. 4j,k, without additional free parameters. The brown line shows a purely passive model in which RNAP does not induce distance-dependent Egr-1 destabilization. The red line shows the experimental passage-time distribution from Fig. 2d. Gray bands represent the simulations intervals containing 90% of simulated datasets. Inset, corresponding mean passage times. Experimental data are shown as mean ± s.e.m.; simulation error bars indicate the central 90% interval of the distribution of replicate means.

However, the model did not fully account for the longest passage times observed in the ZF123 orientation (Fig. S7), which is expected since RNAP-induced destabilization of Egr-1 measured in the static experiments was similar for the two orientations (Fig. 4j,k). This suggests that the larger delay in the ZF123 orientation does not arise solely from TF destabilization and that encounter orientation also affects RNAP dynamics, potentially by increasing the probability of backtrack entry or slowing recovery from backtracked states. Thus, two coupled effects control RNAP passage through Egr-1: orientation-dependent modulation of RNAP translocation dynamics and local destabilization of the TF–DNA complex. This interpretation is consistent with studies of transcription through nucleosomes, where DNA-bound barriers can promote RNAP pausing and backtracking rather than affecting elongation solely through obstacle stability^20,61^.

### CpG methylation modulates Egr-1 binding kinetics and facilitates RNAP passage

Since RNAP passage depends on Egr-1 binding kinetics, we asked whether CpG methylation, a known modulator of Egr-1 dissociation kinetics^50^, also affects passage. We therefore methylated the Egr-1 binding site, which contains two CpG dinucleotides (Fig. S8), and first measured Egr-1 binding kinetics on the methylated constructs. These measurements confirmed that CpG methylation destabilizes Egr-1 binding by increasing its dissociation rate and shortening its residence time on DNA (Fig. S9).

We then repeated the RNAP passage assay using the methylated binding site in the ZF321 orientation (Fig. 6a,b). Consistent with the idea that passage is governed by TF stability, methylation markedly shortened the RNAP passage time in the presence of Egr-1 (Fig. 6c). Indeed, passage through the methylated Egr-1 site was comparable to passage in the absence of TF, indicating that methylation largely eliminates the Egr-1-dependent barrier to elongation. These results show that CpG methylation can regulate RNAP elongation indirectly, by modulating the kinetic stability of DNA-bound transcription factors.

**Figure 6:**
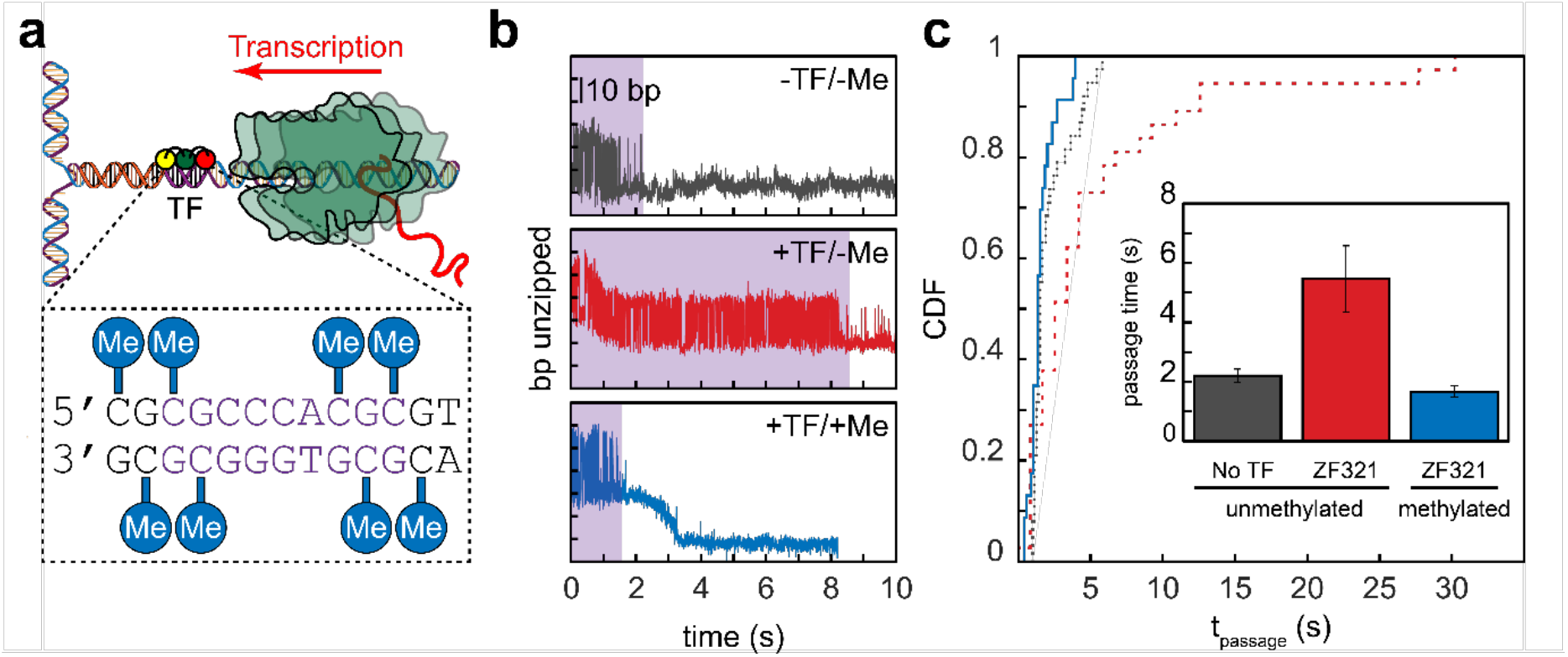
DNA methylation facilitates RNAP passage via Egr-1 destabilization. **a)** Schematic of the RNAP passage assay using a CpG-methylated Egr-1 binding site. The Egr-1 site contains two CpG dinucleotides, which were methylated on both DNA strands. **b)** Representative DNA-fluctuation traces reporting RNAP passage through the fluctuating region in the absence of Egr-1, in the presence of Egr-1 bound to the unmethylated site, and in the presence of Egr-1 bound to the methylated site. RNAP arrival is detected by suppression of DNA fluctuations. **c)** Cumulative distributions of RNAP passage times for the conditions shown in panel b. The distributions for passage in the absence of Egr-1 (dotted line) and for passage through unmethylated Egr-1-bound DNA (dashed line) are reproduced from Fig. 2d and shown here for reference. Statistical comparisons are reported in Table S3. Inset, mean passage times for each condition. Data are mean ± s.e.m.

## DISCUSSION

Transcribing RNA polymerases must frequently navigate DNA-bound proteins, yet the physical mechanisms that allow polymerases to resolve such encounters remain poorly understood. Using Egr-1 as a model DNA-bound transcription factor, we show that RNAP passage through a TF-bound site is governed by a coupled interaction between the elongating polymerase and the bound TF. A bound Egr-1 delays RNAP progression, but RNAP in turn destabilizes the Egr-1–DNA complex. Passage cannot be explained by either of two limiting models: a passive model in which RNAP simply waits for spontaneous TF dissociation, or a mechanical eviction model in which RNAP directly pushes the TF off DNA. Instead, RNAP passage reflects an intermediate mechanism in which TF dissociation, RNAP-induced destabilization, and polymerase backtracking jointly determine the outcome of the encounter.

The intermediate mechanism inferred from the passage experiments implies that RNAP–Egr-1 coupling occurs over a short, spatially structured interaction zone rather than at a single collision point. Static-distance measurements showed that RNAP altered Egr-1 binding kinetics within ~20 bp of the active site, and that this effect was non-monotonic with distance. Surprisingly, Egr-1 could still bind to sites located within the nominal downstream footprint of RNAP, as close as 12 bp from the active site. Structural models place this region of downstream DNA within the RNAP cleft, near jaw, clamp, and lobe elements that engage the downstream duplex^60^. Our ability to detect Egr-1 binding at this position indicates that the RNAP downstream footprint does not represent an absolute steric exclusion zone, but rather a dynamic region in which additional DNA-binding proteins may still bind, albeit with altered kinetics. Incorporating these measured kinetic changes into Monte Carlo simulations accounted for the main features of the passage-time distributions, supporting a model in which local TF destabilization is coupled to RNAP translocation and backtracking.

How might RNAP proximity destabilize Egr-1? One possibility is that contacts between RNAP and downstream DNA locally distort the Egr-1 binding site, weakening zinc-finger contacts without requiring direct protein–protein contact. Elongation complexes bend and clamp downstream DNA, and recent cryo-EM structures of *E. coli* RNAP show that collisions with EcoRI* or with another converging RNAP are accompanied by downstream DNA deformation^62^. Such DNA-mediated allostery^63,64^ could explain why destabilization depends non-monotonically on distance: only specific RNAP–DNA contact geometries would perturb the Egr-1-bound site efficiently. Transient steric or electrostatic contacts between RNAP and Egr-1 may also contribute at short distances. The orientation dependence of passage adds another layer to this mechanism. Although RNAP-induced destabilization of Egr-1 was similar in the two orientations, passage times differed significantly, suggesting that encounter geometry also affects RNAP itself, potentially by increasing the probability of backtrack entry or slowing recovery from a backtracked state. The structural basis of this orientation-dependent effect on RNAP dynamics remains to be determined.

CpG methylation provides a direct example of how the kinetic barrier imposed by a bound TF can be regulated. Consistent with this idea, methylation of the Egr-1 binding site reduced Egr-1 residence time and markedly shortened RNAP passage in the presence of the TF, to a level comparable to passage in the absence of Egr-1. Thus, DNA methylation can modulate transcription elongation indirectly, by changing the kinetic stability of a DNA-bound protein. This observation may be relevant to widespread CpG methylation within mammalian gene bodies^65^. In addition to its known associations with transcriptional activity^65^, CTCF-dependent Pol II pausing^66^, alternative splicing^67^, and suppression of spurious initiation^68^, gene-body methylation may help mitigate TF-dependent kinetic barriers within transcribed regions.

Together, our findings suggest that different DNA-bound obstacles (nucleosomes, high-affinity repressors, and sequence-specific TFs) can impose mechanistically distinct barriers to elongation. For nucleosomes, single-molecule studies support a ratcheting model in which RNAP advances by rectifying transient histone–DNA unwrapping events^20^. High-affinity protein roadblocks such as LacI favor a different regime, involving passive waiting, backtracking, and recovery cycles that can be assisted by transcript-cleavage factors^39,69^. Egr-1 represents another distinct regime: a compact, sequence-specific, moderate-affinity barrier whose effect depends on RNAP-induced kinetic destabilization, motif orientation, CpG methylation, and RNAP backtracking dynamics.

Finally, our results have broader implications for transcription within gene bodies. TF motifs are short and degenerate, making potential binding sites frequent throughout transcribed regions, even outside annotated regulatory elements^24^ (Supplementary Discussion; Fig. S10). Although many of these sites are likely to be occluded by nucleosomes in a given cell type^70^, chromatin accessibility is dynamic and cell-type specific, exposing different subsets of gene-body sequences in different cellular contexts^71^. Such sites need not function as classical regulatory elements to influence transcription: if accessible and bound by a TF, they may create local kinetic barriers to elongating RNAP. Thus, TF binding within gene bodies may provide an additional layer of elongation control, linking chromatin accessibility, TF availability, and transcription dynamics.

## LIMITATIONS OF OUR STUDY

Our study uses a simplified system: bacterial RNAP, naked DNA, the DNA-binding domain of Egr-1, and no accessory elongation factors. This design allowed us to study the mechanism governing RNAP passage through a defined DNA-bound TF. Although elongation factors can facilitate recovery from backtracking, studying the factor-free reaction is informative because RNAPs also possess intrinsic mechanisms for backtrack recovery and transcript cleavage; indeed, yeast cells lacking the TFIIS homolog remain viable, suggesting that intrinsic Pol II activities can be sufficient under many conditions^72^. Nevertheless, this system does not capture the full complexity of eukaryotic transcription through chromatin or the contribution of full-length TF domains. In particular, regulatory disordered regions can modulate TF–DNA binding kinetics and may also introduce additional interactions with RNAP or other elongation-associated factors.

## METHODS

### Proteins

The DNA-binding domain of Egr-1 was expressed and purified as previously described^48^, using a plasmid from Dr. Scot Wolfe, kindly provided by Dr. Amit Meller. *E. coli* RNA polymerase (RNAP) holoenzyme was purchased from NEB (cat. no. M0551S).

### DNA constructs

*DNA handles* were prepared as previously reported^73^, by generating two ~2000-bp DNA segments, each incorporating a specific tag (double digoxigenin and biotin), using commercially purchased 5’-modified primers (IDT; Table S7) in a standard PCR reaction using bacteriophage lambda DNA as the template. The complementary primers (Table S7) were designed to include repeats of three DNA recognition sites for the strand-specific nicking enzymes Nt.BbvCI and Nb.BbvCI (NEB cat. nos. R0632S and R0631S, respectively) on the biotin- and digoxigenin-tagged handles. Nicking generated 29-nt complementary overhangs, allowing the segments to be annealed at equal molar ratios to form a ~4000 bp fragment of connected DNA handles.

A ~280 bp “alignment” segment containing the mouse *Cga* gene promoter sequence was generated via PCR using commercial primers (Table S7) and mouse genomic DNA as a template^74^, then ligated to the annealed handles. The final construct was digested with DraIII-HF (NEB cat. no. R3510L) and kept at −20°C until use.

*Elongation segments* (Table S9) were generated by PCR from pGEM Easy plasmids containing the T7A1 promoter and *LacI* gene. A DraIII restriction site and an Egr-1 binding site were introduced using a partially complementary forward primer (Table S7). A biotin modification was incorporated via the reverse primer for enrichment of transcriptionally active RNAP complexes. PCR products were digested with DraIII-HF (NEB cat. nos. R3510L) following manufacturer protocols.

For experiments probing the effect of methylation, elongation segments were methylated post-amplification and restriction using CpG Methyltransferase (M.SssI, NEB cat. no. M0226L) following the manufacturer’s protocols. To validate methylation, a digestion reaction was performed using the CpG methylation-sensitive HhaI restriction enzyme (NEB cat. no. R0139L). Samples were then analyzed on 1% agarose gels pre-stained with ethidium bromide (Fig. S8).

*AT-rich elongation segments* for RNAP rupture force measurements were prepared by annealing synthetic oligonucleotides (Table S8), as previously described^75^, followed by digestion with DraIII-HF (NEB cat. no. R3510L) per manufacturer instructions.

### Preparation of stalled elongation complexes

10 μL of streptavidin-coated magnetic beads (Invitrogen cat. no. 65606D) were washed on a magnetic separator with 500 μL of washing buffer (20 mM Tris-Cl pH 8.0, 40 mM KCl, 5 mM MgCl_2_, and 1 mM β-mercaptoethanol). To initiate transcription and stall RNAP, the washed beads were incubated with 2.7 μL *E. coli* RNA polymerase holoenzyme (NEB cat. no. M0551S), 1.8 pmol elongation segment, 250 μM ApU dinucleotide (Gena Bioscience cat. no. NU-897L), 50 μM each ATP, GTP, and CTP (NEB cat. no. N0450L), and 0.9 μL RNAsin (NEB cat. no. M0314L), in 35 μL of transcription buffer (25 mM Tris-Cl pH 8.0, 100 mM KCl, 4 mM MgCl_2_, 1 mM DTT, 0.15 mg/mL BSA and 3% v/v glycerol). The reaction was incubated at 37 °C for 15 minutes. After incubation, the beads were separated using a magnetic separator, and the supernatant was removed. The beads were then washed again with 500 μL of ice-cold washing buffer. RNAP complexes were eluted from the beads using AhdI-HF restriction (NEB cat. no. R0584L) for 6 minutes, and stored on ice. (Fig. S1a).

To verify the efficiency of the above procedure in producing elongation competent complexes, 20 μL of streptavidin-coated magnetic beads (Invitrogen cat. no. 65606D) were washed on a magnetic separator with 500 μL of washing buffer. The washed beads were then incubated at 37 °C for 15 minutes in 70 μL of transcription buffer containing 5.4 μL *E. coli* RNA polymerase holoenzyme (NEB cat. no. M0551S), 3.6 pmol of A27-modified T7A1 promoter with an Egr-1 binding site, and 1.8 μL RNAsin (NEB cat. no. M0314L). Following incubation, the beads were separated using a magnetic separator, and the supernatant was removed. The beads were then washed again with 500 μL of ice-cold washing buffer. RNAP-DNA complexes were eluted from the beads by digestion with BglI (NEB cat. no. R0143L) for 15 minutes at 37 °C, following the manufacturer’s protocol with a reduced incubation time. The eluted sample was collected and stored on ice. For stalled complexes and run-off reactions, similar incubation conditions were used with the addition of 250 μM ApU and 50 μM AGC nucleotides. For run-off reactions, stalled complexes were then incubated with 1 mM rNTPs and 36 pmol of a non-biotinylated T7A1 DNA segment for 5 minutes at 37 °C. All reactions were stopped by washing and stored on ice. Elution was performed using AhdI-HF digestion as described. All DNA samples (input, RNAP assembly, stalled complexes, and run-off products) were dialyzed using Slide-A-Lyzer MINI dialysis units (Thermo Scientific cat. no. 69562) in 0.2× TBE buffer for 20 minutes at 4 °C, and analyzed on 1% agarose gels post-stained with ethidium bromide. (Fig. S1b)

### Optical tweezers

Experiments were performed in a custom-made dual-trap optical tweezers apparatus, as previously reported^76,77^. Briefly, the beam from an 852 nm laser (TA PRO, Toptica) is coupled into a polarization-maintaining single-mode optical fiber. The collimated beam out of the fiber is split by a polarizing beam splitter (PBS) into two orthogonal polarizations, each directed into a mirror and combined again with a second PBS. One of the mirrors is mounted on a nanometer scale mirror mount (Nano-MTA, Mad City Labs). A 2x telescope expands the beam and also images the plane of the mirrors into the back focal plane of the focusing microscope objective (Nikon, Plan Apo VC 60x, NA/1.2). Two optical traps are formed at the objective’s focal plane, each by a different polarization, and with a typical stiffness of 0.3-0.5 pN/nm. The light is collected by a second, identical objective, the two polarizations separated by a PBS, and imaged onto two Position Sensitive Detectors (First Sensor). The position of the beads relative to the center of the trap is determined by back focal plane interferometry^78^. Calibration of the setup is done by the analysis of the thermal fluctuations of the trapped beads^79^, which are sampled at 100 kHz. Experiments were conducted using a laminar flow cell (u-Flux, Lumicks), which was passivated following a published protocol^80^ with some modifications. Briefly, the chamber was washed twice by sequentially flushing 1 M NaOH and 1% Liquinox for 10 min each. Casein was sonicated and filtered as a 1% stock solution, diluted to 0.2%, flushed into the chamber, and incubated for 40 min. After incubation, the system was washed with working buffer until free casein was removed.

### Optical tweezers experiments

Elongation segments were ligated to DNA handles using T4 DNA ligase (NEB cat. no. M0202L) and 2× Rapid Ligation Buffer (Promega cat. no. C6711) for 10 minutes at 16 °C. The full construct was incubated for 15 minutes on ice with 0.8 μm polystyrene beads (Spherotech) coated with anti-digoxigenin (anti-DIG). The reaction was then diluted 50-fold in transcription buffer. Tether formation was performed in situ, inside the optical tweezers experimental chamber, by trapping a DNA-bound anti-DIG bead in one trap and a 0.9 μm streptavidin-coated polystyrene bead in the second trap. The beads were brought into close proximity to enable binding of the DNA’s biotin tag to the streptavidin-coated bead. Data were sampled and stored at 2500 Hz. All subsequent processing and analysis were performed using MATLAB. Double-stranded DNA (dsDNA) was modeled using an extensible worm-like chain (eWLC) model, and single-stranded DNA (ssDNA) using a worm-like chain (WLC) model, as previously described^75^. The number of unzipped base pairs was calculated from the measured extension by first subtracting the contribution of the DNA handles, then dividing by twice the extension of a single ssDNA base at the corresponding tension.

*Elongation complex unzipping experiments* were performed by repetitively unzipping DNA constructs harboring an actively elongating RNAP complex. A stalled RNAP was initially assembled and then transferred into a channel containing rNTPs to initiate transcription elongation. The construct was repeatedly unzipped at defined time intervals by moving the steerable trap to introduce an unzipping fork from the upstream end of the template. Each unzipping event produced a force-extension profile in which the position of RNAP along the DNA could be identified as a localized block to fork progression. By aligning successive force traces to the reference DNA map, the position of RNAP was determined with near base-pair resolution. Repeated measurements allowed construction of RNAP position trajectories over time.

*Passage experiments* began by introducing a tethered, stalled RNAP complex into a channel of the laminar flow cell containing 69 nM Egr-1. Several unzipping cycles were performed to verify the initial position of the RNAP and confirm Egr-1 binding. The unzipping fork was then halted after ~290 base pairs, allowing fluctuations in the DNA to be monitored. Next, the flow cell was moved relative to the stationary laser beams to transfer the tether into a second channel containing 0.02–1 mM rNTPs (0.5 mM unless otherwise specified) and 69 nM Egr-1. RNAP passage was detected as a reduction in DNA fluctuations. Control experiments were conducted in the absence of Egr-1.

*RNAP rupture force experiments* were performed using a DNA construct consisting of a T7A1 promoter ligated to a 100% AT-rich segment. In each measurement, a single DNA molecule harboring a stalled RNAP complex was introduced into the optical tweezers chamber and fully unzipped by moving the steerable trap. Upon encountering the bound RNAP, the unzipping fork was blocked and the force monotonically increased. The rupture force was defined as the force at which RNAP dissociated.

*RNAP stall force experiments* were performed using the same experimental geometry as the passage measurements. After confirming the position of the stalled RNAP, the unzipping fork was positioned downstream and the construct was transferred into a channel containing rNTPs to initiate elongation. Data acquisition was continued as RNAP reached and progressively closed the fork. Continued forward translocation shortened the DNA tether and increased the opposing force until RNAP stalled, after which the polymerase backtracked or dissociated. The stall force was defined as the maximum force measured immediately before backtracking or dissociation. Since the RNAP stall-force distributions did not differ significantly among the no-Egr-1, ZF321, and ZF123 conditions (Fig. S4d; Table S5), the measurements were pooled for analysis.

*Egr-1 rupture force experiments* were performed as previously described^48^. A tethered DNA segment containing an Egr-1 binding site was introduced into a channel containing Egr-1, and the steerable trap was moved to unzip the construct. In the presence of a bound protein, progression of the unzipping fork was halted and the applied force increased. The rupture force was defined as the force at which the protein–DNA complex dissociated in a single event. Afterward, the DNA was relaxed to allow re-annealing of the double-stranded DNA.

*Binding-kinetics experiments* were conducted as previously described^50^, with minor modifications. A DNA segment harboring an Egr-1 binding site and a stalled RNAP was unzipped to the position of the Egr-1 site, and the applied force was adjusted to maintain a target probability of the ‘closed’ vs. ‘open’ state. The construct was transferred to a channel containing Egr-1, and fluctuations in extension were recorded. Bound state durations, *t*_*bound*_, were defined as the durations of ‘closed’ states lasting more than 150 ms, while unbound state durations, *t*_*unbound*_, were defined as the time intervals between bound states. The association and dissociation rates (*k*_*on*_and *k*_*off*_) were calculated as the inverse of the mean dwell times, i.e. ⟨*t*_*unbound*_⟩^−1^ and ⟨*t*_*bound*_⟩^−1^, respectively.

### Calculation of Egr-1 binding site density

To identify high-affinity Egr-1 binding sites across gene sequences, we used a position-specific binding energy model based on the single-base substitution data reported by Chattopadhyay et al.^49^. Each 9-bp window along a sequence was scored by computing its relative binding free energy difference (ΔΔG) with respect to the Egr-1 consensus site (GCGTGGGCG), using a position-specific matrix of dissociation constants (KD values) derived from mutational analyses. For sequences with more than three mismatches from the consensus, we applied an additional penalty to account for potential non-additive energetic effects. Predicted K_D_ values were obtained from ΔΔG scores using the relation 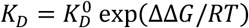, where 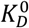 is the dissociation constant for the consensus motif. Sites with *K*_*D*_ < 100 *nM* were classified as high-affinity binding sites. We analyzed the prevalence of such sites in the *Lhb* gene, as well as across a curated set of approximately 3,000 housekeeping genes in the mouse genome. For each gene, the genomic sequence of its coding region was extracted and scanned in both sense and antisense directions using a 9-bp sliding window. We quantified the total number of high-affinity sites, gene length, GC content, and the density of high-affinity sites normalized by gene length.

### Monte Carlo simulations of transcription

The simulation implements a stochastic algorithm to model RNAP passage through a TF binding site. We used a fixed time step (*dt* = 0.01 *s*) approach with non-Markovian dynamics, where transition probabilities depend on both current system state and elapsed times since previous events. RNAP stepping times were drawn from a gamma distribution with shape parameter *α* = 3 and scale parameter θ = (*α* · *k*_*f*_)^−1^, where *k* is the forward stepping rate. Egr-1 association was treated as a pseudo-first-order process and dissociation as a first-order process, with exponentially distributed waiting times. In the absence of RNAP, the baseline rates were 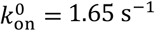 and 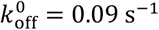, measured for the Δ*x* = 22-bp construct. RNAP-dependent modulation of these rates was implemented separately for association and dissociation using the experimentally measured apparent transition-state energy changes, 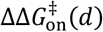 and 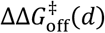 (*d*), where *d* is the RNAP–TF separation. These energy changes were calculated from the measured rates according to

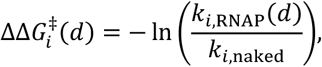

where *i* denotes association or dissociation, and energies are expressed in units of *k*_*B*_*T*. The rates used in the simulations were therefore

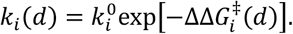

With this convention, positive values of 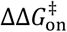 reduce association rates, whereas negative values of 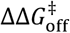 increase dissociation rates. Distance-dependent values of 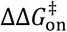 and 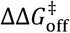 were determined separately for the ZF123 and ZF321 orientations, with linear interpolation between measured distances. For RNAP–TF separations larger than the measured range, 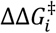 was set to zero, corresponding to unperturbed Egr-1 kinetics. For separations smaller than the minimum measured distance, corresponding to the Δ*x* = 12-bp construct, a very large destabilization factor was applied to reflect steric hindrance. As a control, we also simulated a passive model where RNAP movement is blocked by bound TF but does not alter TF kinetics.

The simulation also incorporates RNAP backtracking, with an exponentially distributed backtrack entry rate *k*_*b*_ and recovery times drawn from the distribution developed by Depken et al.^38^, *f*(*t*) = *t*^−1^ exp(−2*k*_0_*t*) *I*_1_(2*k*_0_*t*), where *I*_1_ is the modified Bessel function of the first kind and *k*_0_ reflects the translocation rate during the backtracked state. Simulations were run until RNAP reached the end position, or a maximum time of 120 *s*. Passage kinetics were analyzed through mean completion times of successful trajectories and their cumulative probability distribution, directly paralleling our experimental measurements.

We inferred the kinetic parameters governing transcriptional elongation (*k*_*f*_) and backtracking (*k*_*b*_ and *k*_0_) from the data in the absence of Egr-1, by fitting an ensemble of simulated passage time distributions to the experimentally measured one. For each candidate parameter set, 10 independent replicates were simulated, each consisting of the same number of trajectories as experimental observations, and the resulting passage time distributions were compared to the experimental data. The empirical cumulative distribution functions (CDFs) were compared using a combined error metric incorporating a weighted mean squared error and the Kolmogorov– Smirnov distance to capture global differences in distribution shape. Parameter optimization was performed using a derivative-free global search algorithm (patternsearch in MATLAB). To estimate 95% confidence intervals for each parameter, we performed nonparametric bootstrapping: the experimental passage times were resampled with replacement, and the optimization procedure was repeated 100 times in parallel. Final confidence intervals were defined by the 2.5th and 97.5th percentiles of the resulting parameter distributions. This procedure resulted in *k*_*f*_ = 47.5 (46.5, 50.3) bp · *s*^−1^, *k*_*b*_ = 1.52 (1.0, 1.96) *s*^−1^, and *k*_0_=1.85 (1.14, 1.95) s^−1^ consistent with previously published values (see for example Mejia et al.^81^).

Using the optimized parameters, the simulations successfully reproduced not only the mean passage times but also the complete distribution of passage times in the presence of TF in the ZF321 orientation. Notably, these predictions contain no additional free parameters, and are based solely on the distribution of passage times in the absence of TF and the binding kinetics in the static experiments, i.e. the results of two other independent experiments, supporting our model.

## ACKNOWLEDGMENTS

We thank Dr. Yael Mandel-Gutfreund and Dr. Dor Shalev for useful discussions. This work was supported by the Israel Science Foundation.

## AUTHOR CONTRIBUTIONS

A.K., P.M., N.N. and S.R. conceived the project. N.N. and S.R. designed all experiments. N.N., S.R. and A.Z. performed the experiments. N.N., S.R., N.S., H.K. and A.K. analyzed the data. C.B. prepared experimental materials. N.N. and A.K. prepared the figures and wrote the manuscript. A.K. supervised the project.

## COMPETING INTERESTS

The authors declare no competing interests.

## Notes

### Competing Interest Statement

The authors have declared no competing interest.

### Summary of Updates

The authors list was updated to reflect the respective contributions

