## Supplementary Material for "Tunable kinetic destabilization governs RNA polymerase passage through a DNA-bound transcription factor"

### Supplementary Information

#### Supplementary Figures

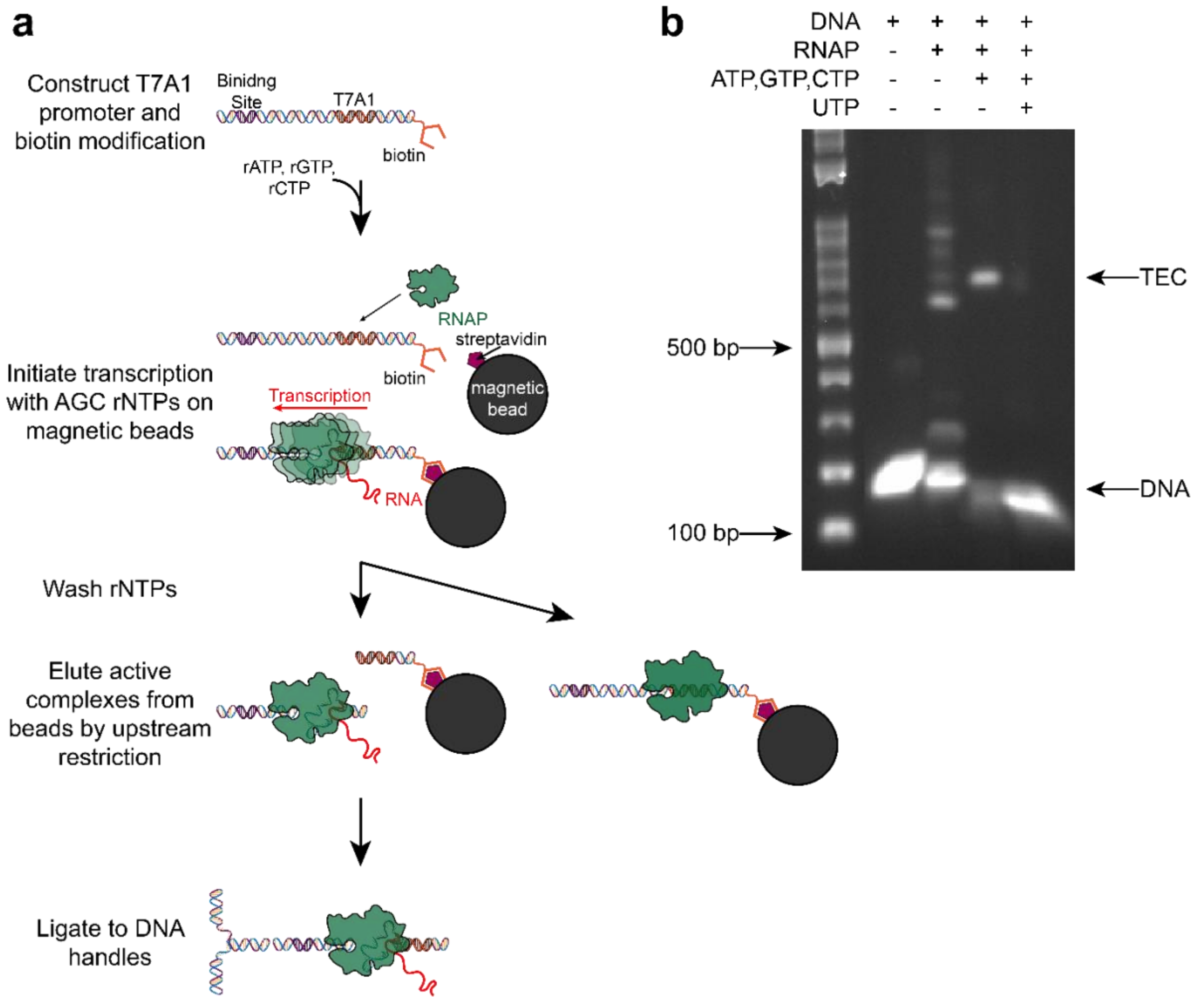


**Figure S1: Assembly and enrichment of elongation-competent stalled complexes. a)** Schematic of the assembly workflow for stalled elongation complexes suitable for detection by DNA unzipping with optical tweezers. In a single reaction, biotin-labeled DNA is bound to streptavidin-coated magnetic beads, RNAP is allowed to form an initiation complex at the promoter, and transcription is initiated in the presence of ApU and three rNTPs: ATP, GTP, and CTP. Omission of UTP causes RNAP to stall after transcription of the 27-bp cassette. The transcription medium and remaining rNTPs are then washed away, and the DNA is digested with AhdI to release elongation-competent complexes from the beads. Because the AhdI site is positioned such that initiation complexes that fail to elongate remain bead-bound, this step enriches the sample for active stalled elongation complexes. The released complexes are then ligated to DNA handles for optical-tweezers measurements. **b)** Gel-based verification of stalled-complex assembly. A 165-bp A27 DNA fragment was incubated with RNAP to verify RNAP association with the DNA. Addition of ApU together with ATP, CTP, and GTP, in the absence of UTP, generated stable stalled transcription elongation complexes. After initiation, complexes were washed and incubated with all four rNTPs, including 1 mM UTP, in the presence of excess decoy DNA to enforce single-turnover conditions. Under these conditions, stalled complexes resumed transcription through the remaining DNA template. Samples were dialyzed and analyzed on a 1% agarose gel post-stained with ethidium bromide.

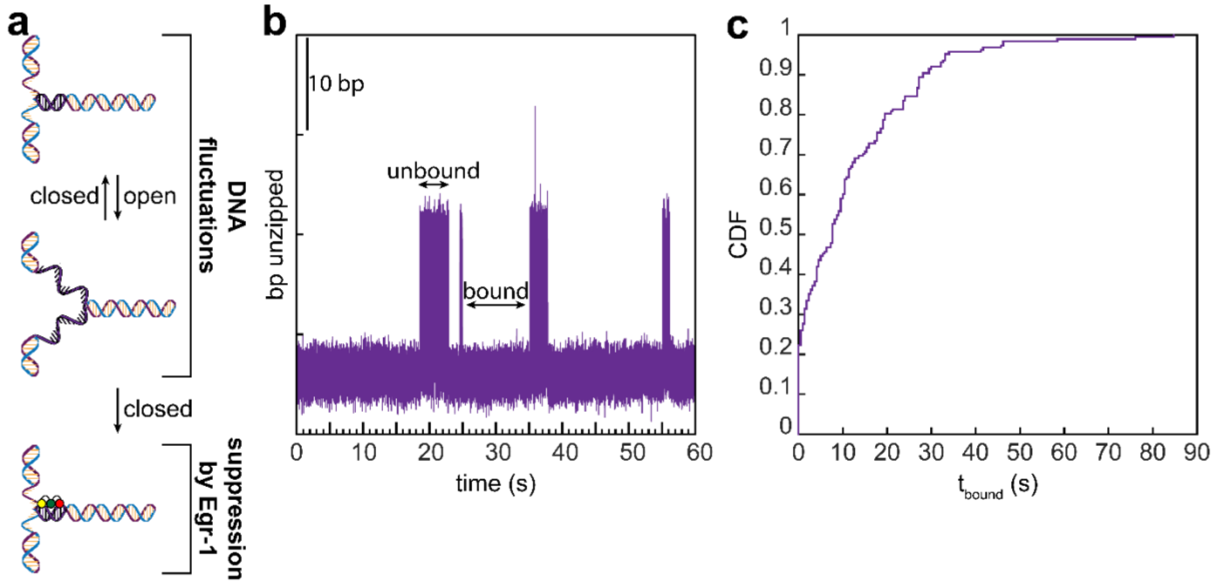


**Figure S2: Residence time of Egr-1 on DNA. a-b)** Schematic (a) and representative trace (b) for the fluctuations assay used to measure the residence time of Egr-1. DNA is partially unzipped to a predetermined position where thermal opening and closing of the DNA fork generate spontaneous fluctuations. Egr-1 binding suppresses DNA fluctuations. **c)** Cumulative distribution of residence times.

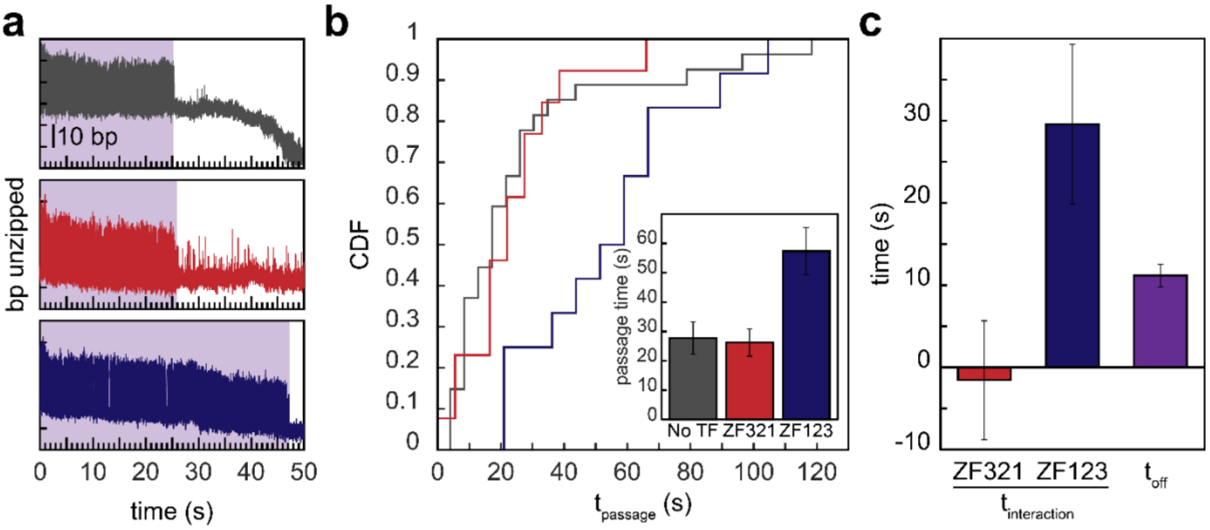


**Figure S3: Egr-1-dependent RNAP passage delay at low rNTP concentration. a)** Representative DNA-fluctuation traces showing RNAP arrival at the fluctuating region under 20 μM rNTP conditions. RNAP passage is faster in the absence of Egr-1 than in the presence of Egr-1 bound in either orientation. Upper, no Egr-1; middle, ZF321 orientation; lower, ZF123 orientation. RNAP arrival is detected by suppression of DNA fluctuations. **b)** Cumulative distributions of RNAP passage times in the absence of Egr-1 and in the presence of Egr-1 in the ZF321 and ZF123 orientations. Inset, mean passage times for the same conditions. Data are mean ± s.e.m.; n=27,13 and 12 molecules for no Egr-1, ZF321, and ZF123, respectively. Statistical comparisons are reported in Table S4. **c)** RNAP–Egr-1 interaction times for the ZF321 and ZF123 orientations, calculated as the difference between the mean passage time measured in the presence of Egr-1 and the mean passage time measured in its absence. Error bars represent propagated s.e.m. from the corresponding passage-time measurements.

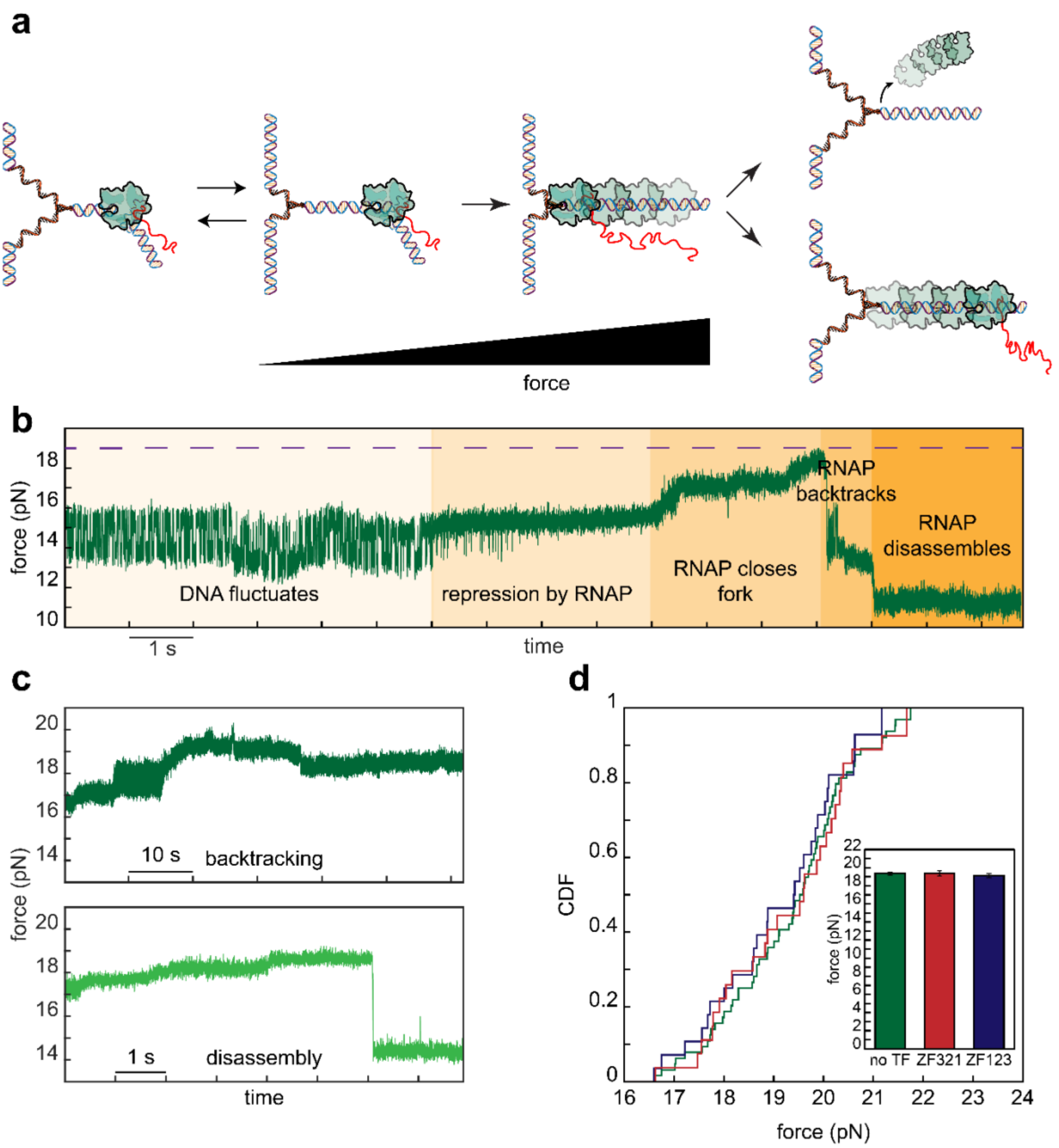


**Figure S4: Measurement of the stall force generated by transcribing RNAP. a)** Schematic of the stall-force assay. The DNA is partially unzipped to generate spontaneous fluctuations between open and closed fork states. Upon RNAP arrival at the fluctuating region, these fluctuations are suppressed. Continued RNAP translocation closes additional segments of the unzipped DNA, increasing the measured force until RNAP stalls and subsequently backtracks or dissociates from the DNA, resulting in a decrease in force. **b)** Representative force trace showing RNAP arrival at the fluctuating region, suppression of DNA fluctuations, and the measured stall force. The magenta dashed line indicates the maximum force. **c)** Representative traces showing two possible outcomes after force buildup. In the upper trace, RNAP backtracks while remaining associated with the DNA and continues to apply force to the fork. In the lower trace, RNAP dissociates from the DNA, causing a sharp decrease in force. **d)** Cumulative distributions of RNAP stall forces measured during transcription in the absence of Egr-1 and after passage through Egr-1 bound in the ZF321 or ZF123 orientation. Inset, mean stall forces for each condition. Data are mean ± s.e.m.; n = 64, 28, and 27 molecules for no Egr-1, ZF321, and ZF123, respectively. Statistical comparisons are reported in Table S5. The stall force was not significantly affected by prior passage through the Egr-1 barrier.

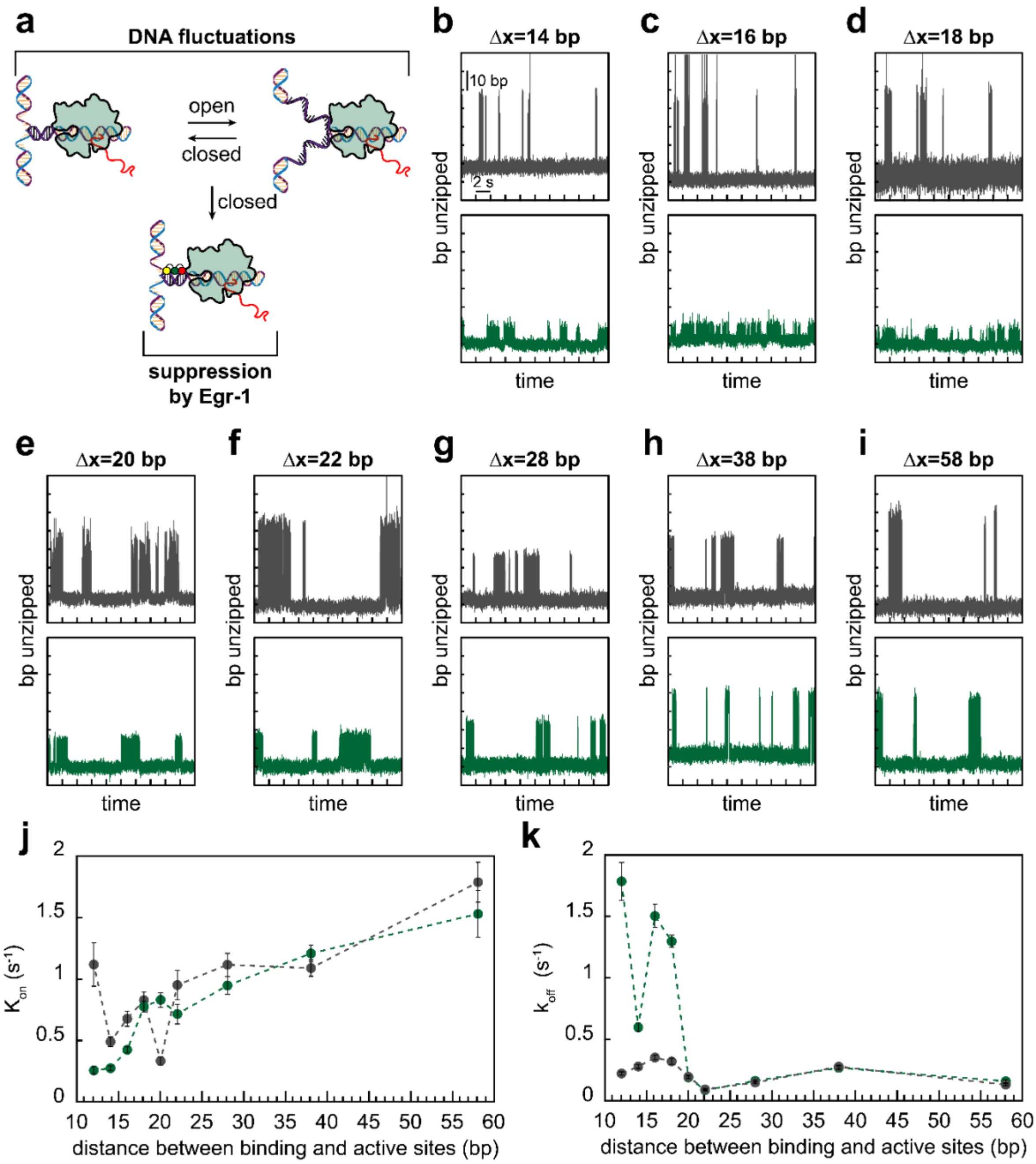


**Figure S5: Effect of RNAP proximity on Egr-1 binding kinetics in the ZF321 orientation. a)** Schematic of the fluctuations assay characterizing Egr-1 binding kinetics near a stalled RNAP complex. Egr-1 binding suppresses fluctuations between the open and closed states of the partially unzipped DNA fork. **b–i)** Representative DNA-fluctuation traces reporting Egr-1 binding and dissociation events in the absence (gray) or presence (green) of RNAP for Egr-1 binding sites located at different distances downstream of the RNAP active site. **j)** Egr-1 association rate, $k_{\text{on}}$, as a function of distance between the Egr-1 binding site and the RNAP active site in the absence and presence of RNAP. **k)** Egr-1 dissociation rate, $k_{\text{off}}$, as a function of distance between the Egr-1 binding site and the RNAP active site in the absence and presence of RNAP. Error bars represent s.e.m.

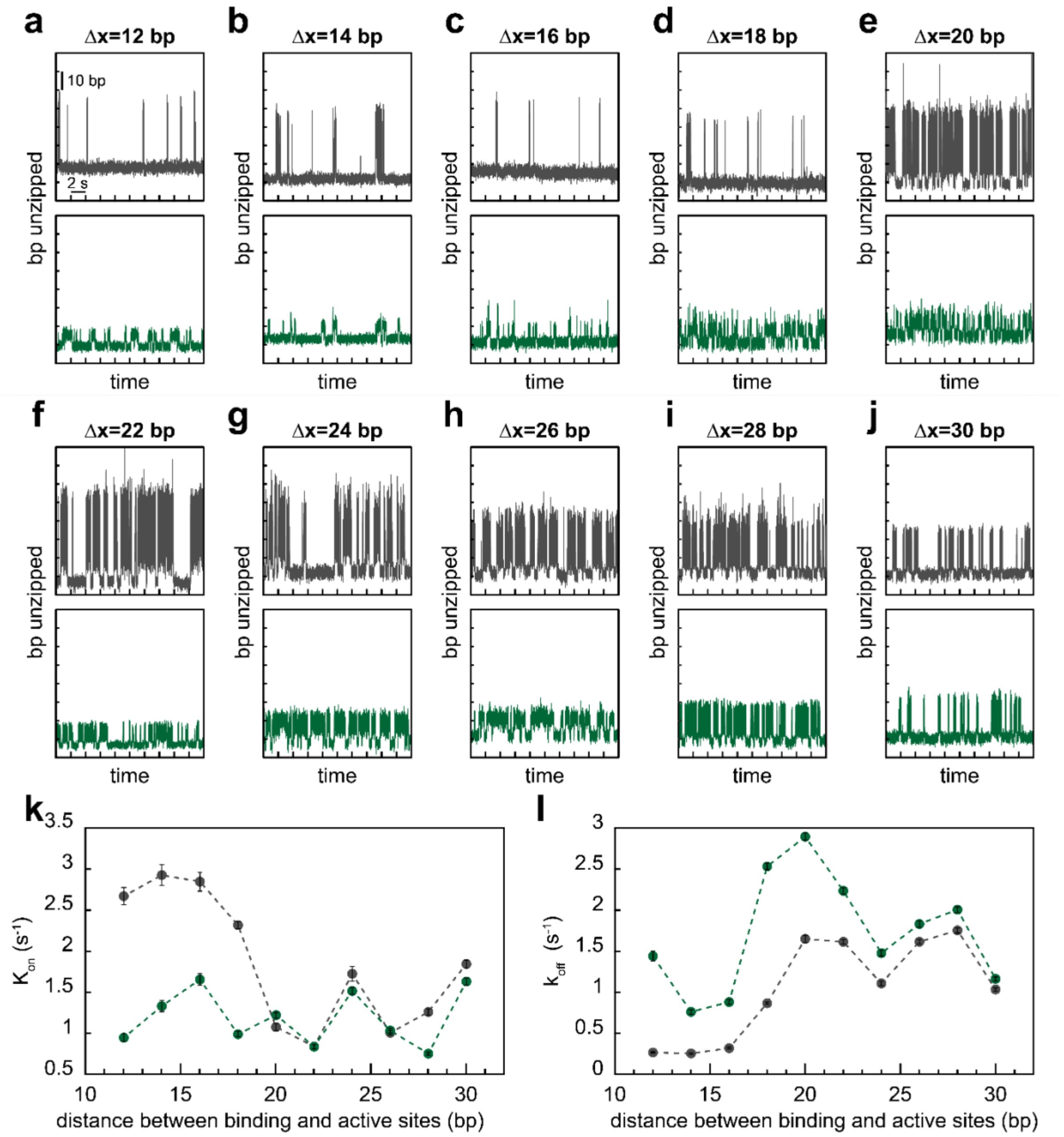


**Figure S6: Effect of RNAP proximity on Egr-1 binding kinetics in the ZF123 orientation. a–j)** Representative DNA-fluctuation traces reporting Egr-1 binding and dissociation events in the absence (gray) or presence (green) of RNAP for Egr-1 binding sites located at different distances downstream of the RNAP active site. **k)** Egr-1 association rate, $k_{\mathrm{on}}$, as a function of distance between the Egr-1 binding site and the RNAP active site in the absence and presence of RNAP. **l)** Egr-1 dissociation rate, $k_{\mathrm{off}}$, as a function of distance between the Egr-1 binding site and the RNAP active site in the absence and presence of RNAP. Error bars represent s.e.m.

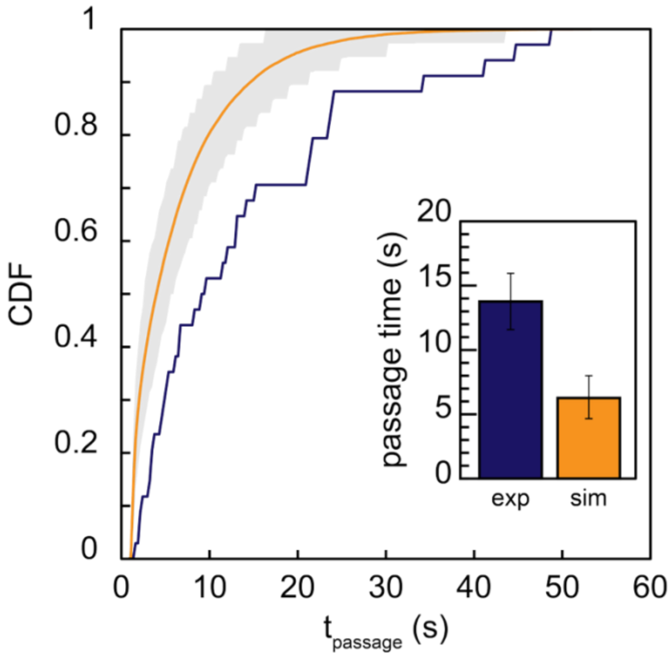


**Figure S7: Simulation results for ZF123 passage.** Stochastic Monte Carlo simulation of RNAP passage in the presence of Egr-1 in the ZF123 orientation. The orange line shows simulations using RNAP elongation parameters obtained from passage on naked DNA (Fig. 5b), and the position-dependent transition-state energy for Egr-1 association and dissociation measured in Fig. 4j,k. The blue line shows the experimental ZF123 passage-time distribution reproduced from Fig. 2d. The gray band represents the simulation interval containing 90% of simulated datasets. Inset, corresponding mean passage times. Experimental data are shown as mean ± s.e.m.; simulation error bars indicate the central 90% interval of the distribution of replicate means.

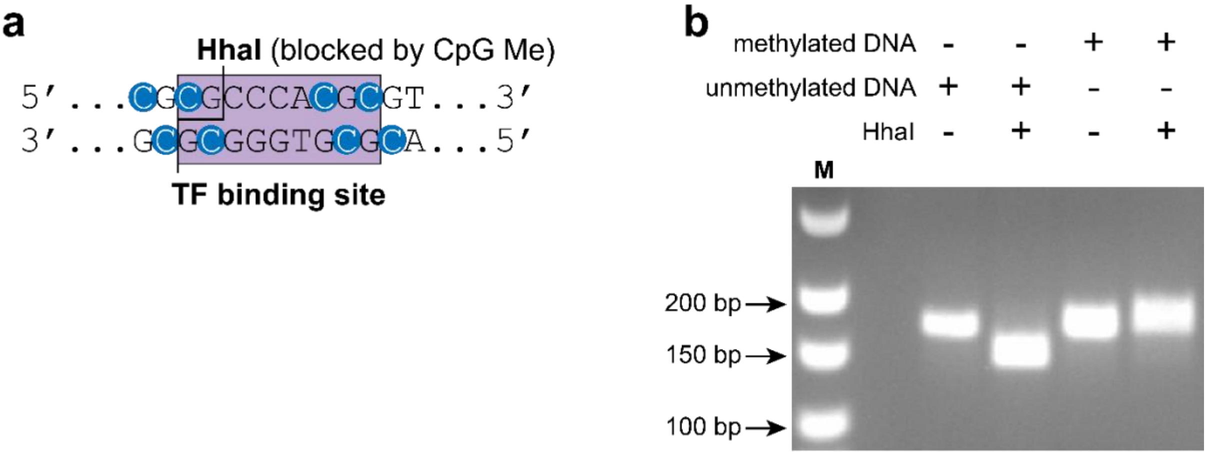


**Figure S8: Verification of CpG methylation within the Egr-1 binding site. a)** Schematic of the Egr-1 binding site, highlighting CpG dinucleotides that overlap the methylation-sensitive HhaI restriction site. CpG methylation blocks HhaI digestion. **b)** Restriction digest verifying CpG methylation. Unmethylated DNA is cleaved by HhaI, resulting in a shorter DNA fragment, whereas methylated DNA is protected from digestion and remains full length. DNA fragments were analyzed by agarose gel electrophoresis.

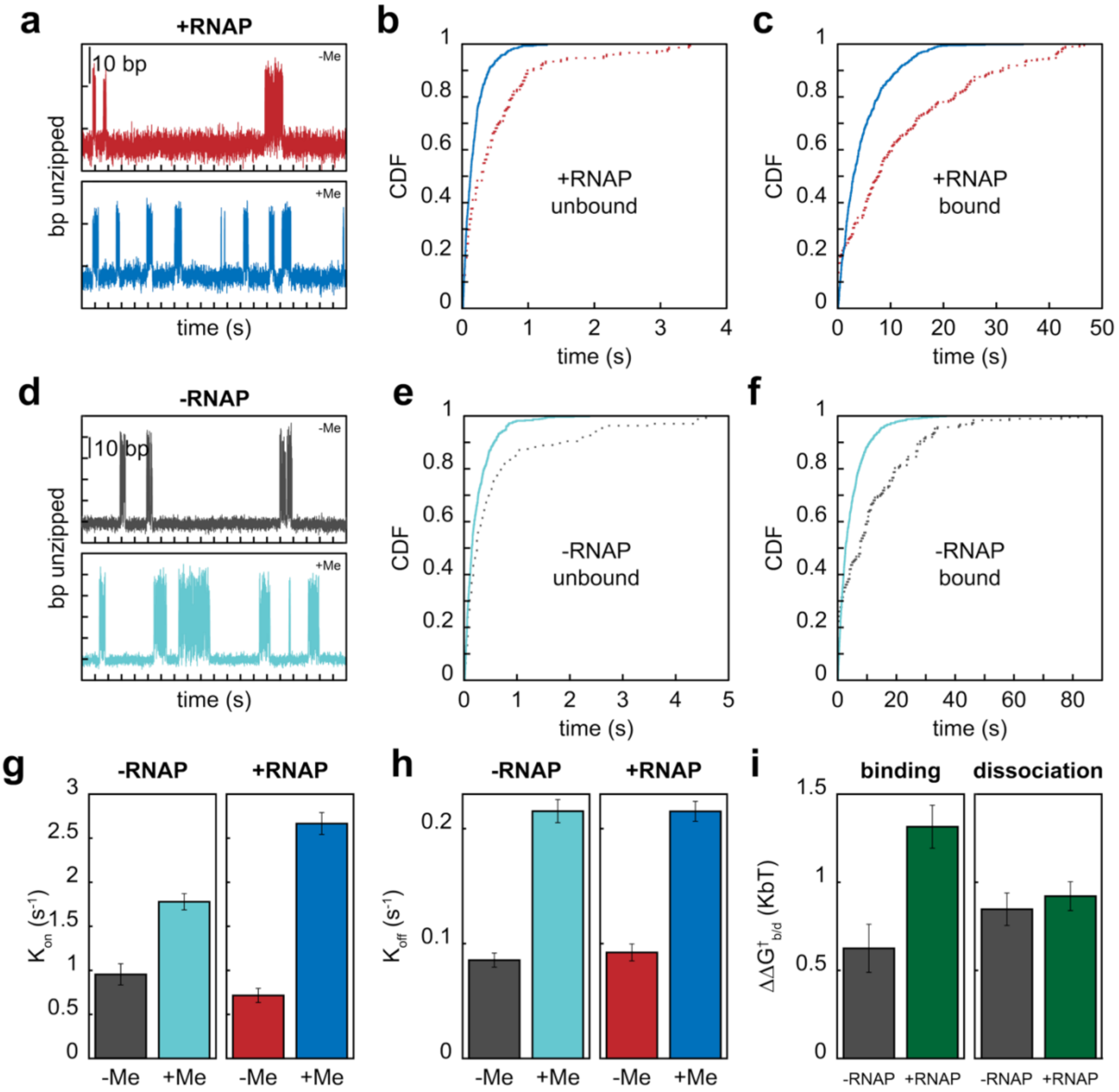


**Figure S9: CpG methylation destabilizes Egr-1 binding to DNA. a)** Representative DNA-fluctuation traces reporting Egr-1 binding and dissociation events in the presence of RNAP on unmethylated DNA (red) and CpG-methylated DNA (dark blue). **b)** Cumulative distributions of unbound-state lifetimes in the presence of RNAP, from which Egr-1 association rates, $k_{\mathrm{on}}$, were extracted. **c)** Cumulative distributions of bound-state lifetimes in the presence of RNAP, from which Egr-1 dissociation rates, $k_{\mathrm{off}}$, were extracted. **d)** Representative DNA-fluctuation traces reporting Egr-1 binding and dissociation events in the absence of RNAP on unmethylated DNA (gray) and CpG-methylated DNA (light blue). **e)** Cumulative distributions of unbound-state lifetimes in the absence of RNAP, from which Egr-1 association rates, $k_{\mathrm{on}}$, were extracted. **f)** Cumulative distributions of bound-state lifetimes in the absence of RNAP, from which Egr-1 dissociation rates, $k_{\mathrm{off}}$, were extracted. **g)** Egr-1 association rates, $k_{\mathrm{on}}$, on unmethylated and methylated DNA in the absence and presence of RNAP. **h)** Egr-1 dissociation rates, $k_{\mathrm{off}}$, on unmethylated and methylated DNA in the absence and presence of RNAP. **i)** Methylation-induced changes in the apparent transition-state energy for Egr-1 association and dissociation. Error bars represent s.e.m. or propagated errors from the corresponding rate estimates, as appropriate.

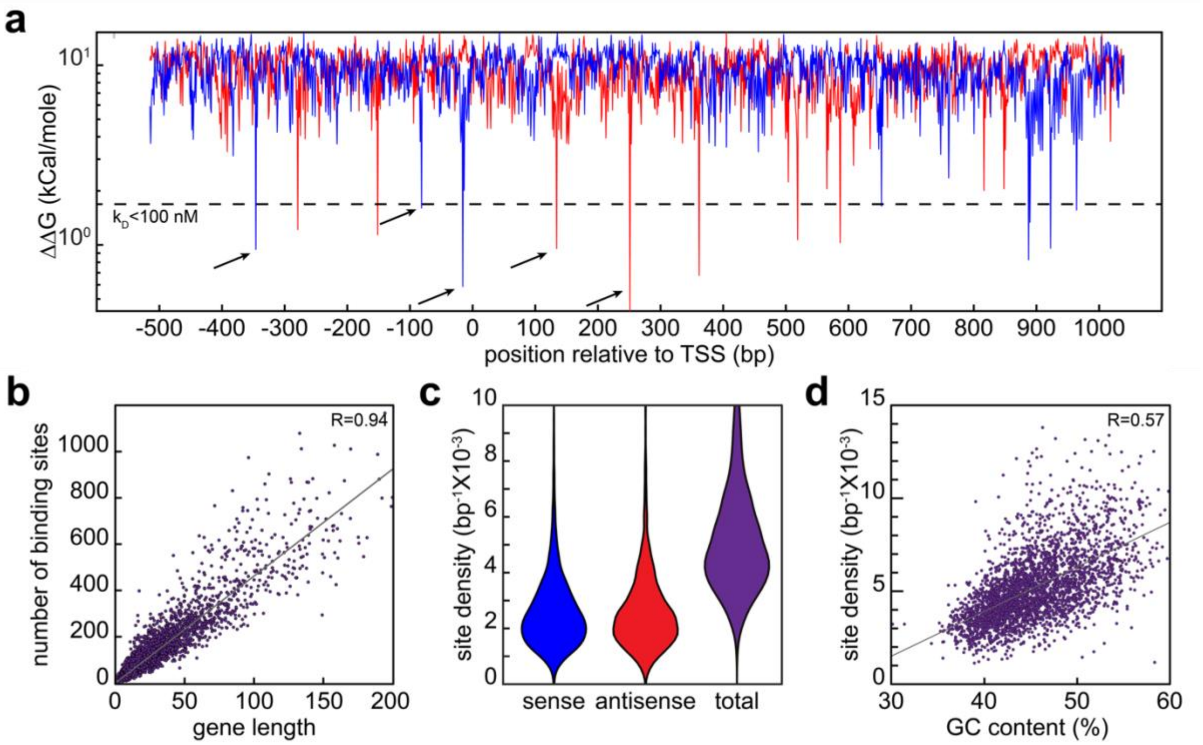


**Figure S10: Egr-1 binding sites are widespread in gene bodies. a)** Predicted binding energy, relative to the consensus Egr-1 binding site, for every 9-bp sequence across the mouse *Lhb* promoter and gene body. Binding energies were calculated for both orientations, shown in blue and red. The dashed line indicates the predicted energy corresponding to $K_D = 100$ nM. Positions below this line were classified as high-affinity Egr-1 binding sites. Predicted high-affinity Egr-1 binding sites are present both in the promoter and within the gene body. **b)** Number of predicted high-affinity Egr-1 binding sites as a function of gene length across the same gene set. For visual clarity, only genes shorter than 200 kbp are shown. Pearson's correlation coefficient (R) is indicated in the plot. **c)** Density of predicted high-affinity Egr-1 binding sites within the gene bodies of ~3,000 mouse housekeeping genes, calculated separately for the sense and antisense orientations and for both orientations combined. Site density was calculated for each gene as the number of predicted high-affinity sites divided by gene length. Statistical comparisons are reported in Table S6. **d)** Predicted high-affinity Egr-1 binding-site density as a function of GC content across the same gene set. For visual clarity, only genes with GC contents between 30% and 60% are shown. Pearson's correlation coefficient (R) is indicated in the plot.

### Supplementary Tables

**Table S1: *p*-values for Fig. 2**
Two-sample Kolmogorov-Smirnov test comparing the distributions of passage times in Fig. 2d.

| | ZF321 (n=37) | ZF123 (n=34) |
|---|---|---|
| No Egr-1 (n=38) | 0.0053 | $5.93 \times 10^{-8}$ |
| ZF321 | | 0.0015 |

**Table S2: p-values for Fig. 3**
Two-sample Kolmogorov-Smirnov test comparing the force distributions shown in Fig. 3g.

| | ZF123 (n=205) | RNAP rupture (n=27) | RNAP stall (n=119) |
|---|---|---|---|
| ZF321 (n=107) | $1.07 \times 10^{-5}$ | $3.59 \times 10^{-10}$ | $1.36 \times 10^{-6}$ |
| ZF123 | | 0.0201 | $2.30 \times 10^{-52}$ |
| RNAP rupture | | | $9.93 \times 10^{-11}$ |

**Table S3: p-values for Fig. 6**
Two-sample Kolmogorov-Smirnov test comparing the distributions of passage times in Fig. 6c.

| | ZF321 (n=37) | methylated ZF321 (n=23) |
|---|---|---|
| No Egr-1 (n=38) | 0.0053 | 0.4641 |
| ZF321 (n=37) | | 0.0012 |

**Table S4: p-values for Fig. S3**
Two-sample Kolmogorov-Smirnov test comparing the distributions of passage times in Fig. S3b.

| | ZF321 (n=13) | ZF123 (n=12) |
|---|---|---|
| No Egr-1 (n=27) | 0.766 | 0.0013 |
| ZF321 | | 0.0145 |

**Table S5: p-values for Fig. S4**
Two-sample Kolmogorov-Smirnov test comparing the distributions of stall forces in Fig. S4d.

| | ZF321 (n=28) | ZF123 (n=27) |
|---|---|---|
| No Egr-1 (n=64) | 0.9946 | 0.9762 |
| ZF321 | | 0.8683 |

**Table S6: p-values for Fig. S10**
Two-sample t-test comparing the site densities in Fig. S10c.

| | antisense | total |
|---|---|---|
| sense | 0.7334 | $<1 \times 10^{-5}$ |
| antisense | | $<1 \times 10^{-5}$ |

**Table S7: Primers for PCR reactions.**

| # | Template | Application | DNA sequence (5' → 3') |
|---|---|---|---|
| 1 | Lambda Phage | Biotin handle | F: /5BiotinTEG/GAT CTC CAG CCA GGA ACT ATT GA<br>R: GTG TCA GCT TGC CCC TCA GCG ATG ACC TCA GCA TTT TTC GAC CTG CTC TTC AGC A |
| 2 | Lambda Phage | DIG handle | F: ATG GCC TAG ACG GCG AGC CTG GGT TTA TAA GGG GAG CGG TGA<br>R: TCA GCT TGC CCC TCA GCG ATG ACC TCA GCA AGG ACC AGC GTT TTG TTG AAA |
| 3 | *Cga* gene promoter | Alignment sequence | F: ATC ACC ACG TGA GAC ATT TTG AGG TAG TGG TG<br>R: ATC ACG CAG TGC TGT TAA TTT AAG AAA TTG GAG CAA TTG T |
| 4 | pGEM easy | -12 stalled RNAP and ZF321 consensus binding site | F: ATC ACT GCG TGG ATG TAT ATA TCT GAC ACG CGC CCA CGC GGC AGT GAC CAT GGT GGT GT<br>R: /5BiotinTEG/ATG GCC TTG CCG GCG TAT AG |
| 5 | pGEM easy | -14 stalled RNAP and ZF321 consensus binding site | F: ATC ACT GCG TGG ATG TAT ATA TCT GAC ACG CGC CCA CGC CCG GCA GTG ACC ATG GTG GT<br>R: /5BiotinTEG/ATG GCC TTG CCG GCG TAT AG |
| 6 | pGEM easy | -16 stalled RNAP and ZF321 consensus binding site | F: ATC ACT GCG TGG ATG TAT ATA TCT GAC ACG CGC CCA CGC CGC CGG CAG TGA CCA TGG TG<br>R: /5BiotinTEG/ATG GCC TTG CCG GCG TAT AG |
| 7 | pGEM easy | -18 stalled RNAP and ZF321 consensus binding site | F: ATC ACT GCG TGG ATG TAT ATA TCT GAC ACG CGC CCA CGC CAC GCC GGC AGT GAC CAT GG<br>R: /5BiotinTEG/ATG GCC TTG CCG GCG TAT AG |
| 8 | pGEM easy | -20 stalled RNAP and ZF321 consensus binding site | F: ATC ACT GCG TGG ATG TAT ATA TCT GAC ACG CGC CCA CGC TTC ACG CCG GCA GTG ACC AT<br>R: /5BiotinTEG/ATG GCC TTG CCG GCG TAT AG |
| 9 | pGEM easy | -22 stalled RNAP and ZF321 consensus binding site | F: ATC ACT GCG TGG ATG TAT ATA TCT GAC ACG CGC CCA CGC GTT TCA CGC CGG CAG TGA CC<br>R: /5BiotinTEG/ATG GCC TTG CCG GCG TAT AG |
| 10 | pGEM easy | -28 stalled RNAP and ZF321 consensus binding site | F: ATC ACT GCG TGG ATG TAT ATA TCT GAC ACG CGC CCA CGC TTA CTG GTT TCA CGC CGG CA<br>R: /5BiotinTEG/ATG GCC TTG CCG GCG TAT AG |
| 11 | pGEM easy | -38 stalled RNAP and ZF321 consensus binding site | F: ATC ACT GCG TGG ATG TAT ATA TCT GAC ACG CGC CCA CGC TCG TAT AAC GTT ACT GGT TTC ACG<br>R: /5BiotinTEG/ATG GCC TTG CCG GCG TAT AG |
| 12 | pGEM easy | -48 stalled RNAP and ZF321 consensus binding site | F: ATC ACT GCG TGG ATG TAT ATA TCT GAC ACG CGC CCA CGC CTC TGC GAC ATC GTA TAA CGT<br>R: /5BiotinTEG/ATG GCC TTG CCG GCG TAT AG |
| 13 | pGEM easy | -58 stalled RNAP and ZF321 consensus binding site | F: ATC ACT GCG TGG ATG TAT ATA TCT GAC ACG CGC CCA CGC CAC CGG CAT ACT CTG CGA CA<br>R: /5BiotinTEG/ATG GCC TTG CCG GCG TAT AG |
| 14 | pGEM easy | -12 stalled RNAP and ZF123 consensus binding site | F: ATC ACT GCG TGG ATG TAT ATA TCT GCT GCC GCG TGG GCG CGT GTT GAC CAT GGT GGT GTT TCC C<br>R: /5BiotinTEG/ATG GCC TTG CCG GCG TAT AG |
| 15 | pGEM easy | -14 stalled RNAP and ZF123 consensus binding site | F: ATC ACT GCG TGG ATG TAT ATA TCT GGC CGG GCG TGG GCG CGT GTA GTG ACC ATG GTG GTG TTT CC<br>R: /5BiotinTEG/ATG GCC TTG CCG GCG TAT AG |

| | | | |
|---|---|---|---|
| 16 | pGEM easy | -16 stalled RNAP and ZF123 consensus binding site | F: ATC ACT GCG TGG ATG TAT ATA TCT GCG GCG GCG TGG GCG CGT GTG CAG TGA CCA TGG TGG TGT TTC C<br>R: /5BiotinTEG/ATG GCC TTG CCG GCG TAT AG |
| 17 | pGEM easy | -18 stalled RNAP and ZF123 consensus binding site | F: ATC ACT GCG TGG ATG TAT ATA TCT GGC GTG GCG TGG GCG CGT GTC GGC AGT GAC CAT GGT GGT<br>R: /5BiotinTEG/ATG GCC TTG CCG GCG TAT AG |
| 18 | pGEM easy | -20 stalled RNAP and ZF123 consensus binding site | F: ATC ACT GCG TGG ATG TAT ATA TCT GGT GAA GCG TGG GCG CGT GTG CCG GCA GTG ACC ATG G<br>R: /5BiotinTEG/ATG GCC TTG CCG GCG TAT AG |
| 19 | pGEM easy | -22 stalled RNAP and ZF123 consensus binding site | F: ATC ACT GCG TGG ATG TAT ATA TCT GGA AAC GCG TGG GCG CGT GTA CGC CGG CAG TGA CCA T<br>R: /5BiotinTEG/ATG GCC TTG CCG GCG TAT AG |
| 20 | pGEM easy | -24 stalled RNAP and ZF123 consensus binding site | F: TCA CTG CGT GGA TGT ATA TAT CTG AAC CAG CGT GGG CGC GTG TTC ACG CCG GCA GTG ACC A<br>R: /5BiotinTEG/ATG GCC TTG CCG GCG TAT AG |
| 21 | pGEM easy | -26 stalled RNAP and ZF123 consensus binding site | F: ATC ACT GCG TGG ATG TAT ATA TCT GGA AAC GCG TGG GCG CGT GGT TTC ACG CCG GCA GTG A<br>R: /5BiotinTEG/ATG GCC TTG CCG GCG TAT AG |
| 22 | pGEM easy | -28 stalled RNAP and ZF123 consensus binding site | F: ATC ACT GCG TGG ATG TAT ATA TCT GGA AAC GCG TGG GCG CGT CTG GTT TCA CGC CGG CAG T<br>R: /5BiotinTEG/ATG GCC TTG CCG GCG TAT AG |
| 23 | pGEM easy | -30 stalled RNAP and ZF123 consensus binding site | F: ATC ACT GCG TGG ATG TAT ATA TCT GGA AAC GCG TGG GCG CGT TAC TGG TTT CAC GCC GG<br>R: /5BiotinTEG/ATG GCC TTG CCG GCG TAT AG |
| 24 | pGEM easy | RNAP structure T7A1 promoter | F: ATG AAG ACT GTA AAT GGT GGT GTT TCC CCG TGT<br>R: /5BiotinTEG/ATG GCC TTG CCG GCG TAT AG |

**Table S8: Oligonucleotides for AT segment construction**

| # | Oligo | DNA sequence (5' → 3') |
|---|---|---|
| 1 | O1 F | ATCACGTCGTGCGGCATGAAGACTATAAATTAATATTAATATTTATATATAAAATTATT TAATATAATTATAAATATTTT |
| 2 | O1 R | TATTTATAATTATATTAAATAATTTTATATATAAATATTAATATTAATTTATAGTCTTC ATGCCGCACGACGTGAT |
| 3 | O2 F | TAAATTATATTTTTATAAATTATTATAAAATTTTTAATTTTAATTATTACATTATTTTA ATATTAAATATAAAAATAA |
| 4 | O2 R | TTTTATATTTAATATTAAAATAATGTAATAATTAAAATTAAAAATTTTATAATAATTTA TAAAAATATAATTTAAAAA |
| 5 | O3 F | AAATTATTTAATTAAATATTTATAAATTATTTTATAATATTATATAAAATAATTTTTTT AATAATTAAAATA |
| 6 | O3 R | TTTATATTTTAATTATTAAAAAAATTATTTTATATAATATTATAAAATAATTTATAAAT ATTTAATTAAATAATTTTTAT |

**Table S9: DNA segments**

| # | Name | DNA sequence (5’ → 3’) |
|---|---|---|
| 1 | Alignment sequence | CACGTGAGACATTTTGAGGTAGTGGTGACCTCAAGGACAGCTTATGAAGAGAGAGCATTTGGCCATTTTCTATGTAATATTATTGACCCTTACACAAAACATCCTGAAAGTTGACACTTTTAAAATAAACAGGACTCTAAGAGTAGCACATTTAATTAGCTAAGTACCTGATATTTTCATAAAGCAGGATAAAAAAAAACATTTTCAGGATTACATTATTTCAACAGGAAACAGAAAAATAAAACAATTGCTCCAATTTCTTAAATTAACAGCACTGC |
| 2 | -12 stalled RNAP and ZF321 consensus binding site | ATCACTGCGTGGATGTATATATCTGACACGCGCCCACGCGGCAGTGACCATGGTGGTGTTTCCCCGTGTCCCTCGATGGCTGTAAGTATCCTATAGGTTAGACTTTAAGTCAATACTCTTTTTGATAATGCGGCCGATGGACCCTATACGCCGGCAAGGCCAT |
| 3 | -14 stalled RNAP and ZF321 consensus binding site | ATCACTGCGTGGATGTATATATCTGACACGCGCCCACGCCCGGCAGTGACCATGGTGGTGTTTCCCCGTGTCCCTCGATGGCTGTAAGTATCCTATAGGTTAGACTTTAAGTCAATACTCTTTTTGATAATGCGGCCGATGGACCCTATACGCCGGCAAGGCCAT |
| 4 | -16 stalled RNAP and ZF321 consensus binding site | ATCACTGCGTGGATGTATATATCTGACACGCGCCCACGCCGCCGGCAGTGACCATGGTGGTGTTTCCCCGTGTCCCTCGATGGCTGTAAGTATCCTATAGGTTAGACTTTAAGTCAATACTCTTTTTGATAATGCGGCCGATGGACCCTATACGCCGGCAAGGCCAT |
| 5 | -18 stalled RNAP and ZF321 consensus binding site | ATCACTGCGTGGATGTATATATCTGACACGCGCCCACGCCACGCCGGCAGTGACCATGGTGGTGTTTCCCCGTGTCCCTCGATGGCTGTAAGTATCCTATAGGTTAGACTTTAAGTCAATACTCTTTTTGATAATGCGGCCGATGGACCCTATACGCCGGCAAGGCCAT |
| 6 | -20 stalled RNAP and ZF321 consensus binding site | ATCACTGCGTGGATGTATATATCTGACACGCGCCCACGCTTCACGCCGGCAGTGACCATGGTGGTGTTTCCCCGTGTCCCTCGATGGCTGTAAGTATCCTATAGGTTAGACTTTAAGTCAATACTCTTTTTGATAATGCGGCCGATGGACCCTATACGCCGGCAAGGCCAT |
| 7 | -22 stalled RNAP and ZF321 consensus binding site | ATCACTGCGTGGATGTATATATCTGACACGCGCCCACGCGTTTCACGCCGGCAGTGACCATGGTGGTGTTTCCCCGTGTCCCTCGATGGCTGTAAGTATCCTATAGGTTAGACTTTAAGTCAATACTCTTTTTGATAATGCGGCCGATGGACCCTATACGCCGGCAAGGCCAT |
| 8 | -28 stalled RNAP and ZF321 consensus binding site | ATCACTGCGTGGATGTATATATCTGACACGCGCCCACGCTTACTGGTTTCACGCCGGCAGTGACCATGGTGGTGTTTCCCCGTGTCCCTCGATGGCTGTAAGTATCCTATAGGTTAGACTTTAAGTCAATACTCTTTTTGATAATGCGGCCGATGGACCCTATACGCCGGCAAGGCCAT |
| 9 | -38 stalled RNAP and ZF321 consensus binding site | ATCACTGCGTGGATGTATATATCTGACACGCGCCCACGCTCGTATAACGTTACTGGTTTCACGCCGGCAGTGACCATGGTGGTGTTTCCCCGTGTCCCTCGATGGCTGTAAGTATCCTATAGGTTAGACTTTAAGTCAATACTCTTTTTGATAATGCGGCCGATGGACCCTATACGCCGGCAAGGCCAT |
| 10 | -48 stalled RNAP and ZF321 consensus binding site | ATCACTGCGTGGATGTATATATCTGACACGCGCCCACGCCTCTGCGACATCGTATAACGTTACTGGTTTCACGCCGGCAGTGACCATGGTGGTGTTTCCCCGTGTCCCTCGATGGCTGTAAGTATCCTATAGGTTAGACTTTAAGTCAATACTCTTTTTGATAATGCGGCCGATGGACCCTATACGCCGGCAAGGCCAT |
| 11 | -58 stalled RNAP and ZF321 consensus binding site | ATCACTGCGTGGATGTATATATCTGACACGCGCCCACGCCACCGGCATACTCTGCGACATCGTATAACGTTACTGGTTTCACGCCGGCAGTGACCATGGTGGTGTTTCCCCGTGTCCCTCGATGGCTGTAAGTATCCTATAGGTTAGACTTTAAGTCAATACTCTTTTTGATAATGCGGCCGATGGACCCTATACGCCGGCAAGGCCAT |
| 12 | -12 stalled RNAP and ZF123 consensus binding site | ATCACTGCGTGGATGTATATATCTGCTGCCGCGTGGGCGCGTGTTGACCATGGTGGTGTTTCCCCGTGTCCCTCGATGGCTGTAAGTATCCTATAGGTTAGACTTTAAGTCAATACTCTTTTTGATAATGCGGCCGATGGACCCTATACGCCGGCAAGGCCAT |

| | | |
|---|---|---|
| 13 | -14 stalled RNAP and ZF123 consensus binding site | ATCACTGCGTGGATGTATATATCTGGCCGGGCGTGGGCGCGTGTAGTGACCATGGTGGTGTTTCCCCGTGTCCCTCGATGGCTGTAAGTATCCTATAGGTTAGACTTTAAGTCAATACTCTTTTTGATAATGCGGCCGATGGACCCTATACGCCGGCAAGGCCAT |
| 14 | -16 stalled RNAP and ZF123 consensus binding site | ATCACTGCGTGGATGTATATATCTGCGGCGGCGTGGGCGCGTGTGCAGTGACCATGGTGGTGTTTCCCCGTGTCCCTCGATGGCTGTAAGTATCCTATAGGTTAGACTTTAAGTCAATACTCTTTTTGATAATGCGGCCGATGGACCCTATACGCCGGCAAGGCCAT |
| 15 | -18 stalled RNAP and ZF123 consensus binding site | ATCACTGCGTGGATGTATATATCTGGCGTGGCGTGGGCGCGTGTCGGCAGTGACCATGGTGGTGTTTCCCCGTGTCCCTCGATGGCTGTAAGTATCCTATAGGTTAGACTTTAAGTCAATACTCTTTTTGATAATGCGGCCGATGGACCCTATACGCCGGCAAGGCCAT |
| 16 | -20 stalled RNAP and ZF123 consensus binding site | ATCACTGCGTGGATGTATATATCTGGTGAAGCGTGGGCGCGTGTGCCGGCAGTGACCATGGTGGTGTTTCCCCGTGTCCCTCGATGGCTGTAAGTATCCTATAGGTTAGACTTTAAGTCAATACTCTTTTTGATAATGCGGCCGATGGACCCTATACGCCGGCAAGGCCAT |
| 17 | -22 stalled RNAP and ZF123 consensus binding site | ATCACTGCGTGGATGTATATATCTGGAAACGCGTGGGCGCGTGTACGCCGGCAGTGACCATGGTGGTGTTTCCCCGTGTCCCTCGATGGCTGTAAGTATCCTATAGGTTAGACTTTAAGTCAATACTCTTTTTGATAATGCGGCCGATGGACCCTATACGCCGGCAAGGCCAT |
| 18 | -24 stalled RNAP and ZF123 consensus binding site | TCACTGCGTGGATGTATATATCTGAACCAGCGTGGGCGCGTGTTCACGCCGGCAGTGACCATGGTGGTGTTTCCCCGTGTCCCTCGATGGCTGTAAGTATCCTATAGGTTAGACTTTAAGTCAATACTCTTTTTGATAATGCGGCCGATGGACCCTATACGCCGGCAAGGCCAT |
| 19 | -26 stalled RNAP and ZF123 consensus binding site | ATCACTGCGTGGATGTATATATCTGGAAACGCGTGGGCGCGTGGTTTCACGCCGGCAGTGACCATGGTGGTGTTTCCCCGTGTCCCTCGATGGCTGTAAGTATCCTATAGGTTAGACTTTAAGTCAATACTCTTTTTGATAATGCGGCCGATGGACCCTATACGCCGGCAAGGCCAT |
| 20 | -28 stalled RNAP and ZF123 consensus binding site | ATCACTGCGTGGATGTATATATCTGGAAACGCGTGGGCGCGTCTGGTTTCACGCCGGCAGTGACCATGGTGGTGTTTCCCCGTGTCCCTCGATGGCTGTAAGTATCCTATAGGTTAGACTTTAAGTCAATACTCTTTTTGATAATGCGGCCGATGGACCCTATACGCCGGCAAGGCCAT |
| 21 | -30 stalled RNAP and ZF123 consensus binding site | ATCACTGCGTGGATGTATATATCTGGAAACGCGTGGGCGCGTTACTGGTTTCACGCCGGCAGTGACCATGGTGGTGTTTCCCCGTGTCCCTCGATGGCTGTAAGTATCCTATAGGTTAGACTTTAAGTCAATACTCTTTTTGATAATGCGGCCGATGGACCCTATACGCCGGCAAGGCCAT |
| 22 | RNAP structure T7A1 promoter | GTGCGGCATGAAGACTATAAATTAATATTAATATTTATATATAAAATTATTTAATATAATTATAAATATTTTTAAATTATATTTTTATAAATTATTATAAAATTTTTAATTTTAATTATTACATTATTTTAATATTAAATATAAAAATAAAAATTATTTAATTAAATATTTATAAATTATTTTATAATATTATATAAAATAATTTTTTTAATAATTAAAATATAAATGGTGGTGTTTCCCCGTGTCCCTCGATGGCTGTAAGTATCCTATAGGTTAGACTTTAAGTCAATACTCTTTTTGATAATGCGGCCGATGGACCCTATACGCCGGCAAGGCCAT |

#### Supplementary Discussion

##### Potential Egr-1 binding sites are frequent within gene bodies

Given the substantial effect of a bound TF on RNAP elongation, we asked how frequently potential TF binding sites occur within gene bodies. We first examined the *Lhb* gene, whose Egr-1 binding sites we previously characterized experimentally[1,2]. Using the previously reported thermodynamic additivity of base-pair substitutions within the Egr-1 binding site[3], we calculated the binding energy, relative to the consensus site, for every 9-bp sequence across the promoter and gene body. Sites with binding energies similar to those of the three known regulatory promoter sites were observed in both orientations (Fig. S10a). Upon stimulation, Egr-1 levels are estimated to reach $\sim 10^4$ copies per nucleus[4,5], corresponding to an effective nuclear concentration of $\sim 150\ nM$, assuming a spherical nucleus with a radius of $\sim 3\ \mu m$. We therefore focused on sites with estimated $K_D \leq 100\ nM$, which are likely to be substantially occupied during periods of Egr-1 activity. Several predicted high-affinity sites were identified within the gene body in both orientations. Notably, the first two gene-body sites in the reverse orientation were previously detected experimentally in single-molecule unzipping experiments and exhibited rupture forces similar to those of the promoter sites.

We next analyzed the density of high-affinity Egr-1 binding sites within the gene bodies of a set of $\sim 3000$ housekeeping genes[6] (Fig. S10b-d). Although the number of sites varied between genes, it correlated strongly with gene length (Fig. S10b), and the overall site density was similar in the sense and antisense orientations (Fig. S10c), and correlated with GC content (Fig. S10d). These trends suggest that the occurrence of high-affinity Egr-1-like sites within gene bodies is largely driven by sequence composition rather than by selection for specific regulatory sites.

Overall, predicted high-affinity Egr-1 sites occurred on average $\sim 2.3$ times per 1000 bp of transcribed DNA in each orientation, corresponding to a mean density of $\sim 2.3 \cdot 10^{-3}$ sites per bp. We next asked how this value compares with a simple random-sequence expectation. The density of any specific 9-bp motif in a sequence scanned with a 1-bp sliding window can be approximated as $4^{-9} = 3.8 \cdot 10^{-6}$ sites per bp in a single orientation, assuming equal nucleotide frequencies and independent positions. However, Egr-1 binding is highly degenerate: of the 2,619 possible single, double, and triple substitution variants relative to the consensus motif, 525 were predicted to bind with $K_D \leq 100\ nM$. The expected density of such high-affinity variants is therefore $3.8 \cdot 10^{-6} \times 525 \approx 2 \cdot 10^{-3}$ sites per bp, in good agreement with the density observed in housekeeping gene bodies. This agreement supports the idea that high-affinity Egr-1-like sites occur frequently within gene bodies as a consequence of motif degeneracy and sequence composition.

Finally, Egr-1 represents only one of many TFs expressed in a given nucleus. The observed density of potential Egr-1 sites therefore suggests that gene bodies, particularly accessible or nucleosome-depleted regions, may contain numerous potential binding sites for DNA-binding proteins. Thus, encounters between elongating RNAPs and bound TFs are expected to be common, highlighting the importance of the passage mechanisms described here.
